# A robust approach for MicroED sample preparation of lipidic cubic phase embedded membrane protein crystals

**DOI:** 10.1101/2022.07.26.501628

**Authors:** Michael W. Martynowycz, Anna Shiriaeva, Max T. B. Clabbers, William J. Nicolas, Sara J. Weaver, Johan Hattne, Tamir Gonen

## Abstract

Crystallization of membrane proteins, such as G protein-coupled receptors (GPCRs), is challenging and frequently requires the use of lipidic cubic phase (LCP) crystallization methods. These typically yield crystals that are too small for synchrotron X-ray crystallography, but ideally suited for the cryogenic electron microscopy (cryoEM) method microcrystal electron diffraction (MicroED). However, the viscous nature of LCP makes sample preparation challenging. The LCP layer is often too thick for transmission electron microscopy (TEM), and crystals buried in LCP cannot be identified topologically using a focused ion-beam and scanning electron microscope (FIB/SEM). Therefore, the LCP needs to either be converted to the sponge phase or entirely removed from the path of the ion-beam to allow identification and milling of these crystals. Unfortunately, conversion of the LCP to sponge phase can also deteriorate the sample. Methods that avoid LCP conversion are needed. Here, we employ a novel approach using an integrated fluorescence light microscope (iFLM) inside of a FIB/SEM to identify fluorescently labelled crystals embedded deep in a thick LCP layer. The crystals are then targeted using fluorescence microscopy and unconverted LCP is removed directly using a plasma focused ion beam (pFIB). To assess the optimal ion source to prepare biological lamellae, we first characterized the four available gas sources on standard crystals of the serine protease, proteinase K. However, lamellae prepared using either argon and xenon produced the highest quality data and structures. Fluorescently labelled crystals of the human adenosine receptor embedded in thick LCP were placed directly onto EM grids without conversion to the sponge phase. Buried microcrystals were identified using iFLM, and deep lamellae were created using the xenon beam. Continuous rotation MicroED data were collected from the exposed crystalline lamella and the structure was determined using a single crystal. This study outlines a robust approach to identifying and milling LCP grown membrane protein crystals for MicroED using single microcrystals, and demonstrates plasma ion-beam milling as a powerful tool for preparing biological lamellae.

## Main

G protein-coupled receptors (GPCRs) are membrane proteins critical to physiological functions in the human body (Lagerström & Schiöth, 2008). Determining GPCR structures using traditional X-ray crystallography is challenging and typically requires crystallization in lipidic cubic phase (LCP) (Landau & Rosenbusch, 1996). Extracting crystals from the viscous LCP is difficult, and many membrane protein crystals only grow to be a few micrometers in size. Structural investigations of GPCRs first turned to X-ray free electron lasers (XFEL) with injector-based LCP delivery systems (Weierstall *et al*., 2014). After years of development, this became a tractable approach. However, XFEL sources are costly, access is highly competitive, and data processing is difficult. In this approach, many individual crystals are typically used during an XFEL experiment, and data from several thousands are then merged to determine a structure (Liu *et al*., 2013). Single particle cryogenic electron microscopy (cryoEM) is an alternative that does not require crystallization, but the small size of most GPCRs prior to the binding of a signaling partner often makes this approach untenable (Liang *et al*.,2017; Danev *et al*., 2021). Microcrystal electron diffraction (MicroED) is a cryoEM method that determines protein structures from single nanocrystals, and is ideally suited to determine these structures (Nannenga & Gonen, 2019). However, the challenges associated with preparing LCP embedded samples for MicroED experiments have thus far limited the use of this method for these critically important structures.

Recent MicroED investigations have reported structures of membrane proteins in viscous media by focused ion-beam milling and subsequent MicroED data collection. In the case of the functional mutant of the murine voltage dependent ion-channel crystallized in lipid bicelles, optimized blotting and dilution on-grid eventually allowed crystal edges to be identified by FIB/SEM (Martynowycz *et al*., 2020). Zhu et al. demonstrated MicroED data collection from LCP embedded crystals of proteinase K by converting the LCP to a less viscous mixture using additives that allowed the liquid layer to be easily blotted away (Zhu *et al*.,2020). However, this approach failed when tested on crystals of membrane proteins. Polovinkin et al. demonstrated diffraction data from bacteriorhodopsin grown in LCP (Polovinkin *et al*., 2020). In this example, a single bacteriorhodopsin crystal over 50 μm wide was looped, placed on an EM grid, milled using a gallium ion-beam, and electron diffraction confirmed the unit cell. However, no structure was determined in this investigation for various technical reasons. We previously determined the structure of the human adenosine receptor A_2A_AR from a single microcrystal (Martynowycz, Shiriaeva *et al*., 2021). To make this sample amenable to MicroED data collection, the crystals were grown in syringes to avoid the rapid dehydration observed from looping crystals from a glass plate and transferring them onto an EM grid. Instead, LCP was converted to the sponge phase inside the syringe. This approach allowed the microcrystal mixture to flow more easily and excess material could be blotted away. Grids were made from this sponge phase mixture and blotted using standard protocols. The microcrystals in the blotted sponge phase grids were visible by FIB/SEM and could be thinned by the gallium beam and subsequently determined by MicroED (Martynowycz & Gonen, 2021). In this work, the looped crystals could be kept hydrated using a humidifier, but resulted in thick layers of ice on the grids, and no crystals could be identified in the FIB/SEM. Thus far, this approach has not been successful for other membrane proteins tested as the conversion to the sponge phase may damage the crystals.

Although these advances have made membrane proteins such as GPCRs more accessible to MicroED, two fundamental issues continue to prevent more widespread adoption: locating crystals in thick media, and making the sample thin enough for MicroED experiments. All reports of membrane structures from milled crystal lamellae have relied on visibly identifying the crystals from topological images in the FIB/SEM. Moreover, conversion of the LCP to the sponge phase may damage the crystals limiting the usefulness of such an approach. To tackle more challenging structures, methods must be developed to successfully mill unconverted LCP and locate crystals inside of the deposited viscous LCP. A fundamental issue with attempting ion-beam milling of LCP embedded crystals is that this material is exceptionally difficult to mill using a gallium beam. Typically, the LCP will begin to indent, turn black, and then deform rather than being removed from the sample. Milling under these conditions is essentially impossible and has prevented milling into thicker LCP areas on the grids. Finding a method to mill away LCP without changing the phase requires a new approach to milling thick samples that does not involve a standard gallium ion beam.

Thinning vitrified biological specimens using a focused ion-beam of gallium ions has become a standard method to prepare samples for electron cryo-tomography (cryoET) and macromolecular microcrystal electron diffraction (MicroED) experiments (Marko *et al*., 2007; Schaffer *et al*., 2017; Martynowycz *et al*.,2019*a*). Unfortunately, milling biological specimens using gallium ions has several drawbacks. For example, gallium sources have limited angular intensity and spherical aberrations that limits their use to relatively low ion-beam currents (Tesch *et al*., 2008). Lower currents increase the amount of time needed to prepare a sample. Furthermore, the gallium ions used for thinning can compromise the experiment by implanting within the sample during milling (Koddenberg *et al*., 2021), and high-energy gallium ions damage the exposed surfaces of the lamellae (Eder *et al*., 2021; Kelley *et al*., 2013). These damaged faces lower achievable signal-to-noise ratio. Thinning biological specimens using a gallium beam is particularly challenging for embedded samples, where only a handful of usable crystal or cellular lamellae are prepared over an entire day (Beale *et al*., 2020) and is slow and inefficient even with automation (Buckley *et al*., 2020; Klumpe *et al*., 2021).

Plasma focused ion beams (pFIBs) are often used in materials science and room-temperature slice-and-view imaging of plastic embedded samples (Gorelick & de Marco, 2019; Binkley *et al*., 2020). Plasma sources are preferable to liquid metal ion-sources for rapid sample preparation, because they maintain coherence at higher beam currents (Smith *et al*., 2006). Additionally, some plasma ion sources, such as xenon, have a higher sputter rate than gallium. This suggests that xenon has the potential to mill faster and cause less radiation damage to the sample than gallium. A recent report using hard materials compared the implantation of ions from various plasma beams after milling tungsten filaments demonstrated that xenon resulted in the lowest implantation depth and shortest milling times, and that oxygen and nitrogen beams lead to oxide and nitride formation within these samples. These reports are in agreement with the stopping range of ions in matter (SRIM) simulations showing that higher Z sources tend to sputter material faster and damage the surfaces less (Eder *et al*., 2021). This approach has not been tested on biological macromolecules but preparing biological lamellae using a pFIB should potentially be faster and increase the signal-to-noise ratio of the subsequently collected data on a TEM. This faster milling along with lower damage might enable creating lamellae of GPCR crystals buried deep within thick, viscous piles of LCP for subsequent MicroED experiments.

Here, we develop methods to create lamellae of vitrified biological material using a plasma focused ion-beam (pFIB) and correlate images in the pFIB/SEM with an integrated fluorescent and light microscope (iFLM) on-the-fly. First, we characterize the four available plasma ion sources - xenon, argon, nitrogen, or oxygen - to prepare lamellae of vitrified biological samples at cryogenic temperatures. To quantitatively assess the outcomes, we vitrified microcrystals of the serine protease, proteinase K, on EM grids. Microcrystals were machined for each ion source using the same protocols to a target thickness of 300 nm. This roughly corresponds to the inelastic mean free path of electrons accelerated through 300 kV and typically leads to the highest quality data (Martynowycz, Clabbers *et al*., 2021). Grids with milled lamellae were transferred into a cryogenically cooled transmission electron microscope (TEM), and continuous rotation MicroED (Nannenga *et al*., 2014) data were collected in electron counting mode on a direct electron detector (Martynowycz *et al*., 2022). The data quality, images, quantitative and qualitative features between the various ion-beam sources were compared between individual lamellae. Structures were determined from data from each gas source to compare the quality of the resulting models. Next, we prepared frozen grids containing fluorescently labelled human adenosine receptor containing BRIL fusion protein in the third intracellular loop and a C-terminal truncation of residues 317 to 412 (A_2A_AR-BRIL-ΔC, hereafter A_2A_AR) in LCP. While crystals were not visible under the thick LCP layer by SEM, they were clearly visible by fluorescence allowing efficient targeting for milling. Crystals were identified deep within thick piles of LCP using fluorescent microscopy and correlated to images taken by the SEM to precisely target the crystals. Fluorescent images were taken periodically while lamellae were prepared deep within the sample using the plasma ion beam. The structure of A_2A_AR was then determined by MicroED using a single milled crystal to a resolution of 2.7 Å.

### A plasma focused ion-beam (pFIB) for vitrified biological specimens and targeting using fluorescence

A Helios Hydra 5 CX dual-beam (Thermo-Fisher) instrument equipped with a cryogenically cooled stage was employed for these investigations. This instrument allows for the selection of either xenon, argon, oxygen, or nitrogen ion sources to form a pFIB and an improved SEM column compared to instruments used in prior investigations (**Methods**) (Skalicky *et al*., 2016; Martynowycz *et al*., 2019*a*,*b*). The sample stage operates at a 4 mm working distance that roughly corresponds to the coincidence point between the electron and ion beams that are oriented 52° apart. A new sample shuttle for this system holds two clipped TEM grids at a pre-tilt of 27°. The system has an integrated fluorescence light microscope (iFLM) that operates with a 20 × objective with an imaging field of view of approximately 350 μm with a working distance of approximately 600 μm. The light microscope operates in either reflective or fluorescence mode using one of four selectable excitation wavelengths (λ= 385, 470, 565, 625 nm). Light microscopy is conducted by translating the sample within the chamber and rotating the shuttle 180° from the standard imaging and milling orientation. Integration of the light microscope allows for on-the-fly identification of targets and correlative light and electron microscopy (CLEM). The integrated light microscope is designed based on the photon ion electron microscope (PIE scope) as described (Gorelick *et al*., 2019). Tagging the protein with a fluorophore prior to crystallization enables the unambiguous identification of protein crystals embedded in thick material. In this way, proteins that are buried in thick media can be identified that would otherwise be impossible by other means.

We hypothesized that the iFLM could be used to target fluorescently labelled GPCR crystals that were buried in LCP on an EM grid. Protein of A_2A_AR was fluorescently labelled prior to crystallization. The crystals grown in LCP were then spread on an EM grid using a crystallography loop and frozen in liquid nitrogen. Screening these grids, no crystals were visible topologically using either the SEM or pFIB beams (**Figure 1A**). However, translating the stage to the iFLM allowed immediate identification of crystals buried under the surface of the LCP piles (**Figure 1B**). Upon identification, a stack of images was taken in Z to target the crystal location (where it is the most in focus and the maximum fluorescent signal is). Overlaying the fluorescence data onto the SEM images allowed pinpointing the crystal coordinates in the pFIB/SEM (**Figure 1C, D**). However, several obstacles prevented accurately milling deep into the LCP. Namely, the SEM and iFLM data needed to be correlated to the grazing incidence milling pFIB beam, the samples need to be protected from the powerful plasma ion-beam, and the best plasma ion-beam for obtaining the highest quality data had to be determined.

**Figure 1.**
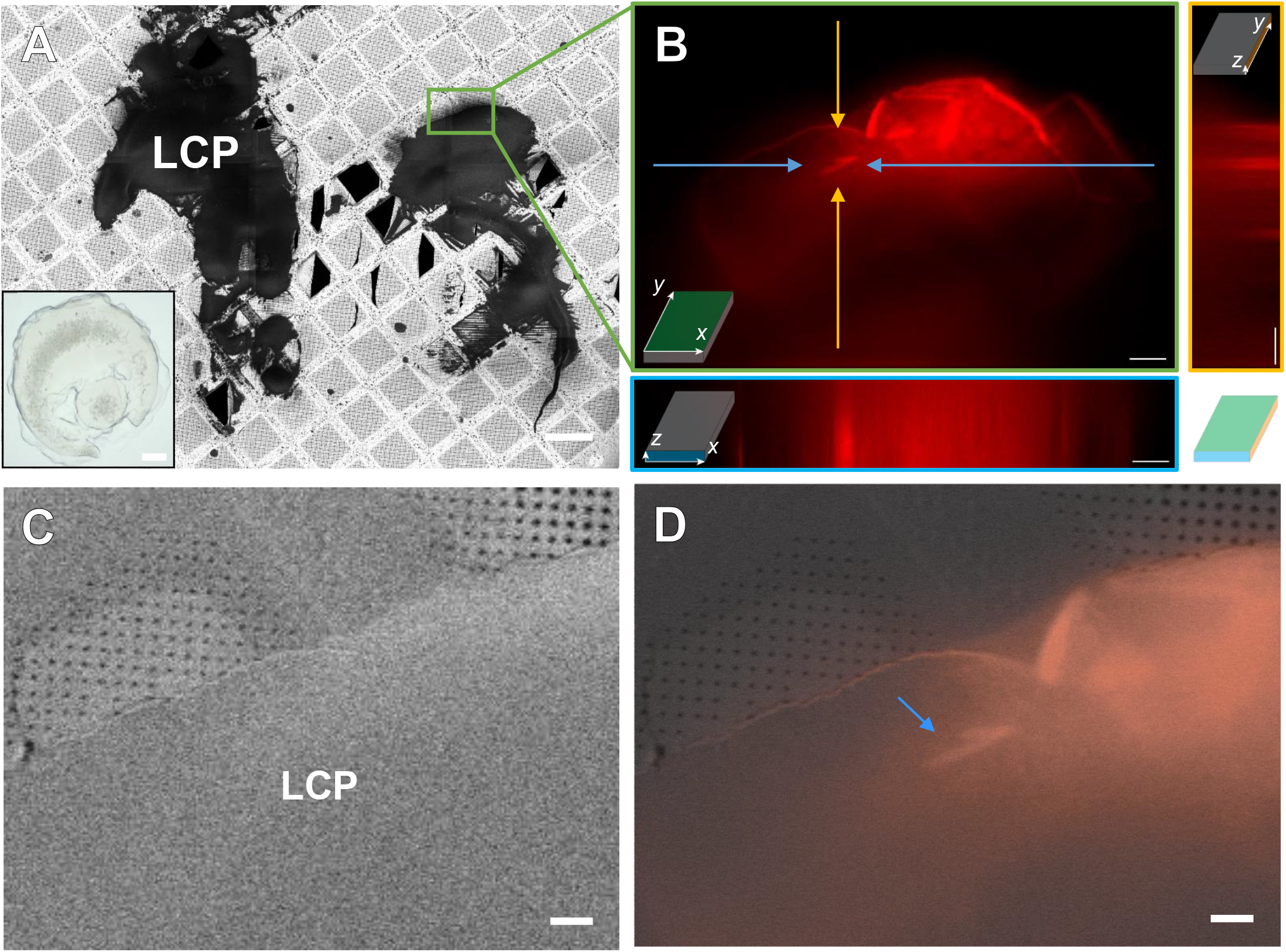
Identification and targeting of fluorescently labelled GPCR microcrystals in thick LCP. (A) 500 V SEM image montage of a prepared grid of A_2A_AR in LCP. Inset shows a typical A_2A_AR drop. Scale bar 100 μm. (B) Fluorescent image from a Z-stack taken from the location highlighted from (A) in the green box. The bottom and right panels depict the projection of the stack in either the Z-X or Y-Z planes, respectively. Arrows to the crystal are color coded to their locations in the corresponding projections. Scale bars 25 μm. (C) SEM image of the area in (B) taken after platinum coating. Scale bar 10 μm. (D) Correlative overlay of (B) onto (C) showing the location of the GPCR crystals deep in the LCP. Scale bar 10 μm.

Because the crystals were buried deeply in the thick LCP, we had to determine a way to target them as accurately as possible in the Z-dimension. X- and Y-dimensions are relatively accurate but the Z-dimension (depth) resolution is relatively poor in brightfield cryogenic fluorescent light microscopy. For this we first calibrated the iFLM using fluorescent beads (4 micron Tetraspecs) embedded in a thick matrix of 50% glycerol to mimick the viscosity of the LCP. We alternated between milling and imaging to correlate the iFLM measured depth of the beads and the disappearance depth of the beads measured by the angled view of the pFIB. (**Supplementary Figure 1**, **Methods**). A similar approach was described previously for cryoET applications (Arnold *et al*., 2016). Using this method we were able to reliably target regions of interest buried deep in thick media.

Even at low flux, the ion-beam can damage the sample during imaging and milling (Zhou *et al*., 2019). Milling is typically conducted at much higher beam currents than imaging (Schaffer *et al*., 2017; Beale *et al*.,2020; Martynowycz & Gonen, 2021). Although milling is contained to a defined region, the beam is usually much larger than the defined area milled. The spilled over exposures build up at the sample face over the course of the experiment. Additionally, making lamellae with an even thickness requires the front of the sample to be nearly homogenous and smooth. For this purpose, a layer of platinum was deposited to protect the samples using the gas injection system (GIS) at a grazing incidence modified for this use case (**SI Figure 2, Methods**). With the milling depth estimation and GIS protection strategy sorted, it was necessary to fully characterize the pFIB sources for milling vitrified biological material.

### Thinning biological samples using different plasma sources

Microcrystals of a serine protease, proteinase K, were grown in batch and vitrified onto TEM grids. The grids were placed into autogrid clips and loaded into the pFIB/SEM and coated in GIS platinum. Twenty crystals were identified on a single grid using SEM imaging (**Figure 2A**). Five crystals were milled using each ion source-20 crystals in total (**Figure 2, SI Figs 3–6**). Each crystal was milled at approximately 15°, corresponding to a stage tilt of 4° with an 11° sample pre-tilt and SEM imaging angle of approximately 67° (**Figure 2**). The milling was conducted using pre-defined cleaning cross sections (**Methods, SI Table 1**). Each lamellae was prepared in four steps to a final target thickness of 300 nm (**Figure 2, SI Table 1**). This thickness is roughly the inelastic mean free path of an electron accelerated through a potential of 300 kV, and was previously determined to maximize MicroED data quality (Martynowycz, Clabbers *et al*., 2021). Each set of five lamellae was milled sequentially. The source gas was then switched, the plasma ion-beam aligned, and the next lamellae were milled. All twenty lamellae across all four gas sources were prepared in a single 10-hour shift.

**Figure 2.**
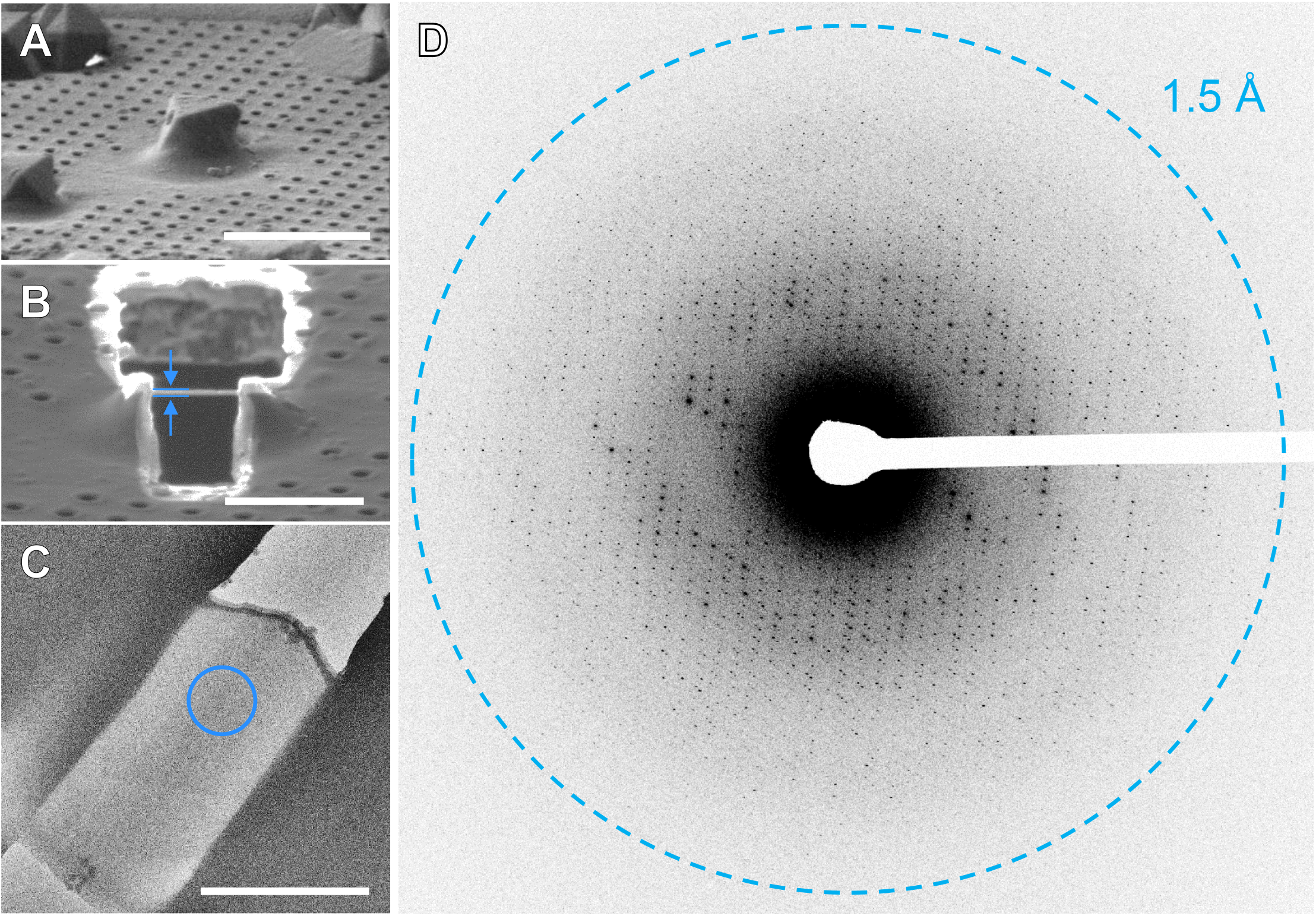
Preparing plasma beam milled lamellae of a protein microcrystal. Images of a selected serine proteinase microcrystal before (A) and after (B) thinning the crystal into a thin lamella using a focused ion-beam of argon ions. This lamella showing clear delineation of the platinum layer, crystal, and vitrified media at 2200 × in the TEM (C). (D) MicroED data corresponding to 4° of data summed together from a direct electron detector.

**Table 1.** MicroED structures of Proteinase K determined from plasma beam milled lamellae

**Figure 5**
|  |  |  |  |  |  |  |  |  |  |
| --- | --- | --- | --- | --- | --- | --- | --- | --- | --- |
| Plasma beam | argon |  | xenon |  | nitrogen |  | oxygen |  | Best Merge |
| Acceleration Voltage (kV) | 300 |  | 300 |  | 300 |  | 300 |  | 300 |
| Wavelength | 0.0197 |  | 0.0197 |  | 0.0197 |  | 0.0197 |  | 0.0197 |
| Resolution range | 19.77 - 1.4 (1.45 - 1.4) |  | 19.74 - 1.45 (1.50 - 1.45) |  | 19.75 - 1.8 (1.86 - 1.8) |  | 20.36 - 1.5 (1.55 - 1.5) |  | 20.63 - 1.39 (1.44 - 1.39) |
| Space group | P 43 21 2 |  | P 43 21 2 |  | P 43 21 2 |  | P 43 21 2 |  | P 43 21 2 |
| Unit cell | 67.02 67.02 107.53<br>90 90 90 |  | 67.05 67.05 107.02<br>90 90 90 |  | 67.12 67.12 106.87<br>90 90 90 |  | 67.26 67.26 106.81<br>90 90 90 |  | 67.02 67.02 107.53<br>90 90 90 |
| Total reflections | 1231493 (96739) |  | 925857 (66300) |  | 354396 (34224) |  | 529482 (41429) |  | 2726169 (150232) |
| Unique reflections | 47738 (4616) |  | 43468 (4250) |  | 23288 (2268) |  | 38542 (3742) |  | 49781 (4770) |
| Multiplicity | 25.8 (20.7) |  | 21.3 (15.5) |  | 15.2 (15.0) |  | 13.7 (11.0) |  | 54.8 (31.3) |
| Completeness (%) | 97.27 (96.17) |  | 98.61 (98.29) |  | 99.68 (99.21) |  | 96.20 (95.51) |  | 99.44 (97.05) |
| Mean I/sigma(I) | 7.61 (1.40) |  | 7.73 (1.62) |  | 4.18 (1.24) |  | 4.91 (1.03) |  | 10.78 (1.68) |
| Wilson B-factor | 11.9 |  | 12.65 |  | 18.53 |  | 9.69 |  | 11.71 |
| R-merge | 0.3235 (1.784) |  | 0.2942 (1.475) |  | 0.5183 (1.798) |  | 0.759 (1.674) |  | 0.3396 (1.773) |
| R-meas | 0.3301 (1.829) |  | 0.3008 (1.525) |  | 0.5374 (1.862) |  | 0.7844 (1.757) |  | 0.3426 (1.802) |
| R-pim | 0.06426 (0.3945) |  | 0.0608 (0.3758) |  | 0.1379 (0.4721) |  | 0.1944 (0.5264) |  | 0.04454 (0.3109) |
| CC1/2 | 0.993 (0.266) |  | 0.995 (0.306) |  | 0.964 (0.299) |  | 0.877 (0.274) |  | 0.997 (0.327) |
| Reflections used in refinement | 47632 (4616) |  | 43393 (4250) |  | 23271 (2268) |  | 38471 (3742) |  | 49735 (4770) |
| Reflections used for R-free | 2329 (242) |  | 2167 (207) |  | 1142 (101) |  | 1965 (211) |  | 2476 (227) |
| R-work | 0.1374 (0.2720) |  | 0.1387 (0.2869) |  | 0.1679 (0.2844) |  | 0.1634 (0.2737) |  | 0.1192 (0.2780) |
| R-free | 0.1735 (0.3108) |  | 0.1770 (0.3488) |  | 0.2121 (0.3796) |  | 0.2138 (0.3431) |  | 0.1634 (0.2964) |
| macromolecules | 2063 |  | 2052 |  | 2031 |  | 2047 |  | 2031 |
| ligands | 10 |  | 10 |  | 2 |  | 10 |  | 6 |
| solvent | 307 |  | 294 |  | 237 |  | 344 |  | 322 |
| Protein residues | 279 |  | 279 |  | 279 |  | 279 |  | 279 |
| RMS(bonds) | 0.015 |  | 0.009 |  | 0.004 |  | 0.002 |  | 0.016 |
| RMS(angles) | 1.1 |  | 0.88 |  | 0.63 |  | 0.48 |  | 1.84 |
| Ramachandran favored (%) | 97.11 |  | 97.47 |  | 96.39 |  | 97.47 |  | 96.75 |
| Ramachandran allowed (%) | 2.89 |  | 2.53 |  | 3.61 |  | 2.53 |  | 3.25 |
| Ramachandran outliers (%) | 0 |  | 0 |  | 0 |  | 0 |  | 0 |
| Rotamer outliers (%) | 0.91 |  | 0 |  | 0 |  | 0 |  | 0 |
| Clashscore | 2.47 |  | 2.24 |  | 5.3 |  | 1.99 |  | 3.02 |
| Average B-factor | 14.47 |  | 15.14 |  | 19.42 |  | 12.16 |  | 14.77 |
| macromolecules | 12.51 |  | 13.43 |  | 18.47 |  | 10 |  | 12.77 |
| ligands | 27.44 |  | 25.25 |  | 20.14 |  | 24.69 |  | 20.47 |
| solvent | 27.21 |  | 26.76 |  | 27.49 |  | 24.63 |  | 27.29 |

We found differences between the gas sources and categorize the qualities of each gas by the following criterion: milling speed, imaging quality, and success rate (**Figure 3, SI Table 1, SI Figures 3 – 6**). By inspection using the ion and electron beams, we found that several crystal lamellae had some signs of cracking, splitting, or being otherwise destroyed during the milling process, most notably nitrogen (5/5) and oxygen (4/5) displayed the most damage to the crystals (**SI Figures 3 – 6**).

**Figure 3.**
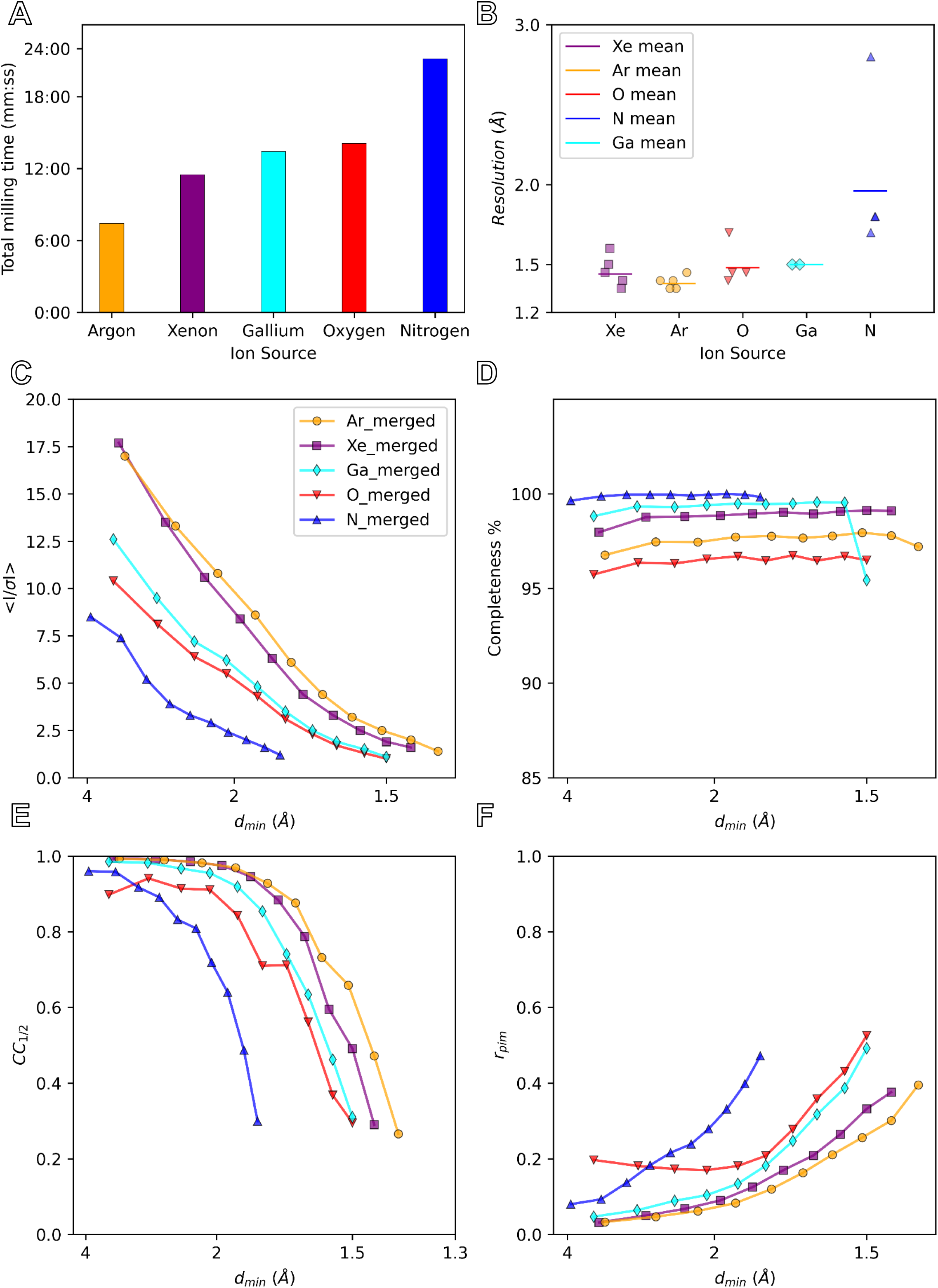
Crystallographic statistics for plasma ion-beam milled lamellae from different sources. Plots depict the total milling time (A), MicroED resolution (B), mean signal to noise ratio (<I / σ (I)>) (C), completeness (%) (D), mean half-set correlation coefficient (CC_1/2_) (E), and merged multiplicity corrected R factor (R_pim_)(F) as functions of the d_min_ resolution bins (Å). The merged datasets are solid lines with symbols with xenon in purple, argon in orange, oxygen in red, nitrogen in blue, and gallium in teal.

Imaging specimens with the plasma ion-beams is similar to using a gallium ion-beam instrument. However, the depth of field was different for each ion source. Adjusting the ion-beam image for any of the plasma sources was more challenging than for gallium sources. The contrast of the images roughly correlates to the mass of the ion—xenon had the best contrast, whereas nitrogen had the worst (**SI Figures 3 – 6**). However, the faster sputter rates for xenon and argon typically made tasks such as focusing the image more challenging, because the area used to focus would rapidly deteriorate at higher beam currents. The oxygen and nitrogen sources have additional blurring due to how the magnetic lenses affect these lighter elements, resulting in ‘double images’ in both the left-right and up-down directions. The left-right double image can be corrected via direct alignments inside the column. However, the top-down double image could not, and was instead corrected by sticking rare earth magnets to the plasma beam column until sharp images could be obtained (**Methods**). Lamellae were transferred into a cryogenically cooled TEM for further investigation (**Figure 2C, D**).

### MicroED data collection

After cryo-transfer into the TEM, we assessed each lamella by visual inspection of low-dose images taken on a direct electron detector (**Figure 2C, SI Figures 7 – 10**). Ice contaminations and breakage not observed in the SEM prior to loading in the TEM are attributed to the cryo transfer step. All 20 lamellae sites were identified in the TEM using low magnification imaging. At higher magnifications, breaks on the far side of 2/5 (1 minor, 1 large) argon milled lamellae became visible along the edges (**SI Figure 8**). Visual inspection of the unbroken or cracked portions of the milled lamellae was used to assess the degree of curtaining on the surface of each crystal using TEM imaging. In this assessment, all xenon lamellae had evidence of strong curtaining and streaks, most oxygen lamellae had visible curtaining that was less severe than xenon, and argon had the least visible curtaining that we could assess (**SI Figures 7 – 10**). The lamellae milled by nitrogen all contained serious visible pathologies, including a hole through the top of the lamella (**SI Figure 9**).

Continuous rotation MicroED datasets were collected identically from each lamella in electron counting mode on a Falcon 4 direct electron detector (**Figure 3F, SI Figures 6–9**) (Martynowycz *et al*., 2022). Data were collected from each lamella using the same rotation rate over an identical real-space wedge (**Methods**). Data were isolated from a 2 μm diameter area using a selected area aperture. In this way, we were able to collect data from nearly all the lamellae (18/20). Maximum intensity projections were calculated to visually inspect the resolution of each dataset prior to processing, since single frames in counting mode contain very little visible signal (**SI Figures 7 – 10**). Electron counting movies were converted to crystallographic format, and then indexed and integrated identically (**Methods**).

### Data quality from different plasma sources

Crystallographic intensity statistics were determined after applying a high-resolution cutoff for each dataset where the mean half-set correlation coefficient (CC_1/2_) fell to approximately 30% (**Figure 3, SI Figures 11–14**) (Karplus & Diederichs, 2012). Data from each ion source were merged to separate averaged results from individual trends (**Figure 3, SI Figures 11 – 14**). In terms of crystallographic statistics, we found that the highest average (< I / σ (I) >) came from lamellae prepared using the argon beam, followed by xenon, oxygen, and nitrogen (**Figure 3**). Completeness was relatively high for each crystal. We attribute differences in completeness to variations in crystal orientation on the grid. The mean half-set correlation coefficient (CC_1/2_) and the redundancy corrected merging R factor, R_pim_, showed the same overall trends as (< I / σ (I) >), where the best results seemingly came from argon, followed by xenon, oxygen, and then nitrogen (**Figure 3**). The statistics from oxygen most closely resemble the best results using gallium ions to mill this protein, whereas both argon and xenon data appear to consistently yield better data.

### Protein structures from plasma milled lamellae

Structures of proteinase K were successfully determined from the merged data of each gas source by molecular replacement (**Figure 4, Table 1, Methods**) (McCoy *et al*., 2007). Each structure was refined using the same settings, with calcium and nitrate ions being added manually when found between refinement cycles (Kovalevskiy *et al*., 2018; Emsley & Cowtan, 2004). The resolutions of the lamellae were 1.40, 1.45, 1.50, and 1.80 Å for argon, xenon, oxygen, and nitrogen, respectively (**Figure 4**). This is compared to our prior best result of 1.5 Å using a gallium ion-beam. After the final rounds of refinement, the R-work and R-free for the same experiments were found to be: 13.74 / 17.35, 13.87 / 17.70, 16.79 / 21.21, 16.34 / 21.38. Surprisingly, the R factors for both argon and xenon milled lamellae were both significantly better than any prior investigation of this protein by MicroED, whereas the R factors for both nitrogen and oxygen were overall similar to those in prior investigations at similar resolutions. The prior best gallium milled structure resulted in an R-work and R-free of 14.95 and 20.46, respectively (Martynowycz *et al*., 2022). The structures determined from plasma milled lamellae all showed well defined side chains and essentially undamaged disulfide bonds (**Figure 3**)(Hattne *et al*., 2018). As expected, the higher resolution structures of xenon and argon show more resolved waters than the lower resolution model derived from nitrogen milled lamellae. Merging across different ion sources was also explored, and the increased multiplicity resulted in even better structural model to compare against the individual merged sets (“Best Merge”, **Figure 2, Table 1**, **SI Figure 15**, **Methods**). The model derived from oxygen milled lamellae, however, showed a significantly larger number of water molecules than expected based on resolution and overall poorer crystallographic refinement, with statistics similar to the lower-resolution model from the nitrogen beam.

**Figure 4.**
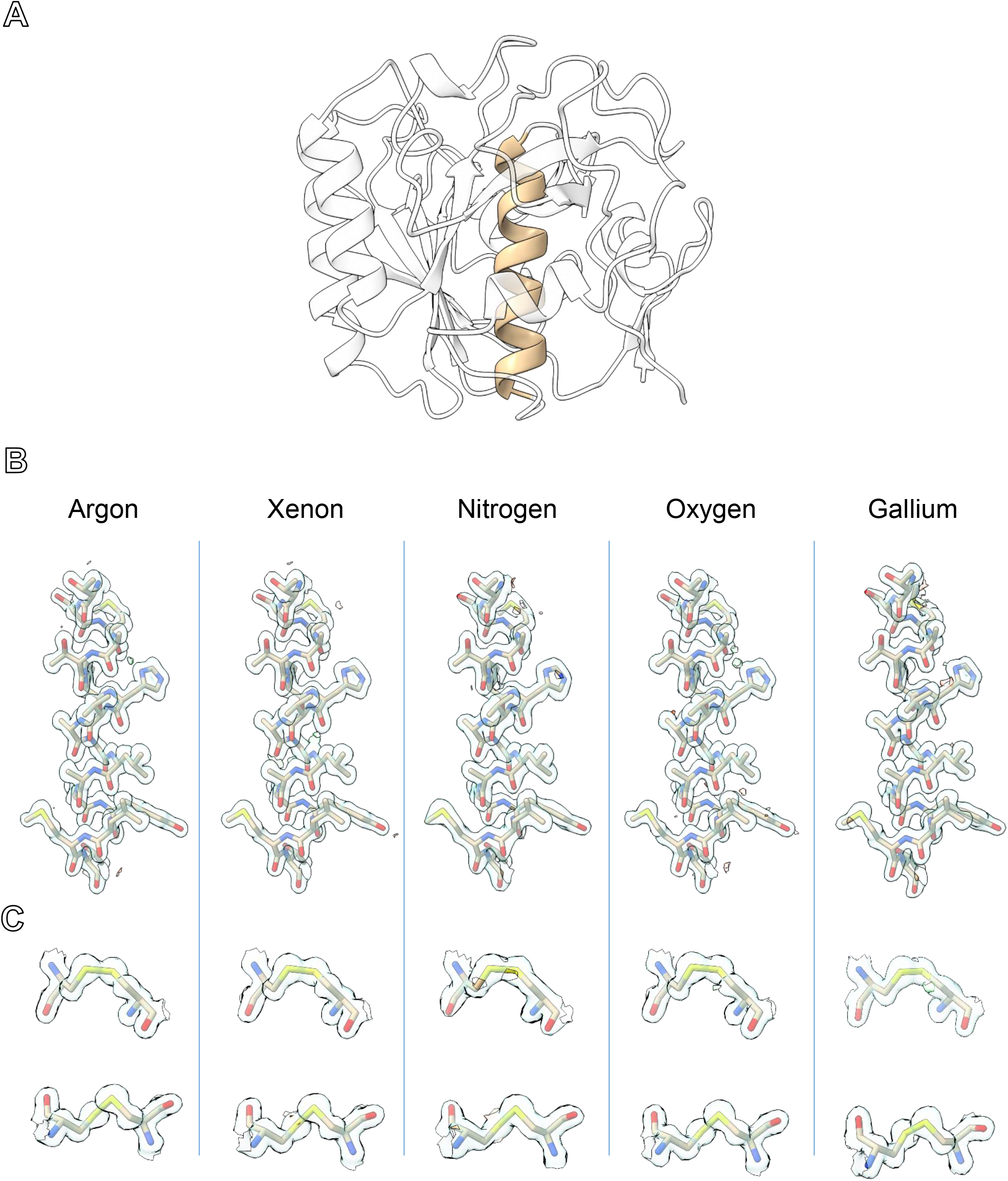
The structure of proteinase K determined from plasma ion-beam milled lamellae. (A) The structure of the serine protease, proteinase K, determined by MicroED from plasma ion-beam milled lamellae. (B) Maps for each plasma source and the prior best gallium structure from the same helix (residues 328 – 344) highlighted in (A). (C) The two disulfide bonds in proteinase K (Cys^139^ – Cys^228^ top, and Cys^283^ – Cys^364^ bottom) for each structure. 2mF_o_-DF_c_ maps are all contoured at the 1.5 σ level, and the mFo-DFc difference maps are all contoured at ± 3 σ level in green and red, respectively.

### Targeting buried GPCR crystals by correlated light and electron microscopy

Grids containing fluorescently labelled A_2A_AR were prepared by looping large amounts of material from crystallization drops in a glass-sandwich plates (**Methods**). To prevent the rapid degradation of these crystals, the looping was done at high humidity. A 100 μm nylon crystallography loop was used to scoop up a large amount of both LCP and A_2A_AR microcrystals, gently scraped along the surface of a pre-clipped EM grid, and immediately plunged into liquid nitrogen (**Methods**). These grids were loaded into the pFIB/SEM at cryogenic conditions. An all-grid atlas was taken of the grid using the SEM at an accelerating voltage of 500 V to better target future positions and increase contrast (**Figure 1A**). The grid was then coated in a protective layer of platinum using the GIS similarly to the grids containing proteinase K. From the overview, areas of LCP were identified that were between 5 and 100 μm above the holey carbon film. No crystals were visible from any angle using either the SEM or pFIB (**Figure 1C**). Instead, the stage was translated and inspected using the iFLM using either the reflective mode, where no filter cube was used, or by one of the four wavelengths. Crystals could not be identified using the reflective mode imaging, but the latter was useful to evaluate the topology of the sample and estimate the height of the surface. However, the red and green fluorescent channels successfully identified crystals buried deep within the LCP (**Figure 1B**). By taking multiple images over a range of focal distances, we were able to identify the depth of the crystal relative to the surface of the LCP and from the position of the underlying grid bars below. The fluorescent and reflective stacks were simultaneously correlated to the X-Y plane of the SEM images (**Figure 1D**). In this way, the position of an A_2A_AR crystal was determined in three dimensions to enable targeting of essentially invisible crystals buried in the thick LCP (**Methods**). The crystal selected for milling was approximately 20 μm above the holey carbon film and approximately 10 μm from the top of the LCP. Additional crystals were nearby in this same pile, but were all directly over a grid bar, rendering them unusable (**Figure 1D**).

A lamella was created from the selected A_2A_AR crystal using the xenon beam. Xenon was chosen because of its high sputter rate for these extremely deep sites. Although the argon beam was faster for the small serine proteinase crystals, this was limited by the breaking of the crystals rather than the sputter rate of the ion. Due to the immense size of the LCP occluding the crystal, initial milling was conducted at 15 nA, a current that would not be possible for milling frozen samples in a gallium ion system. The current was stepped down as the lamella approached the physical crystal location in Z (**Methods**). Between each thinning step, one or more fluorescent images were taken at the crystal focal plane to assure the crystal was not destroyed or over-milled. The final lamella was approximately 10 μm wide, 250 nm thick, and required the removal of at least 10 × 40 × 50 μm of LCP, carbon, and ice from either above or below the suspended crystal. The plasma ion beam showed no deformation or decoloring of the LCP (**Figure 5A**). Due to the increased current and sputter rate, the total milling time on this lamellae was under 1 hr, with the majority of time for the experiment being taken by imaging and checking the sample between milling steps. A final stack of fluorescent images was taken from the thin lamellae and correlated to an SEM image and confirmed the crystal survived the milling process (**Figure 5B**). From the fluorescent image taken at the focal plane of the lamella, we could see the milled crystal appeared sharper in the lamella than the unmilled portion outside of the lamella, indicative of the higher noise from the LCP that was not removed from this area (**Figure 5B**,blue versus white arrow).

**Figure 5.**
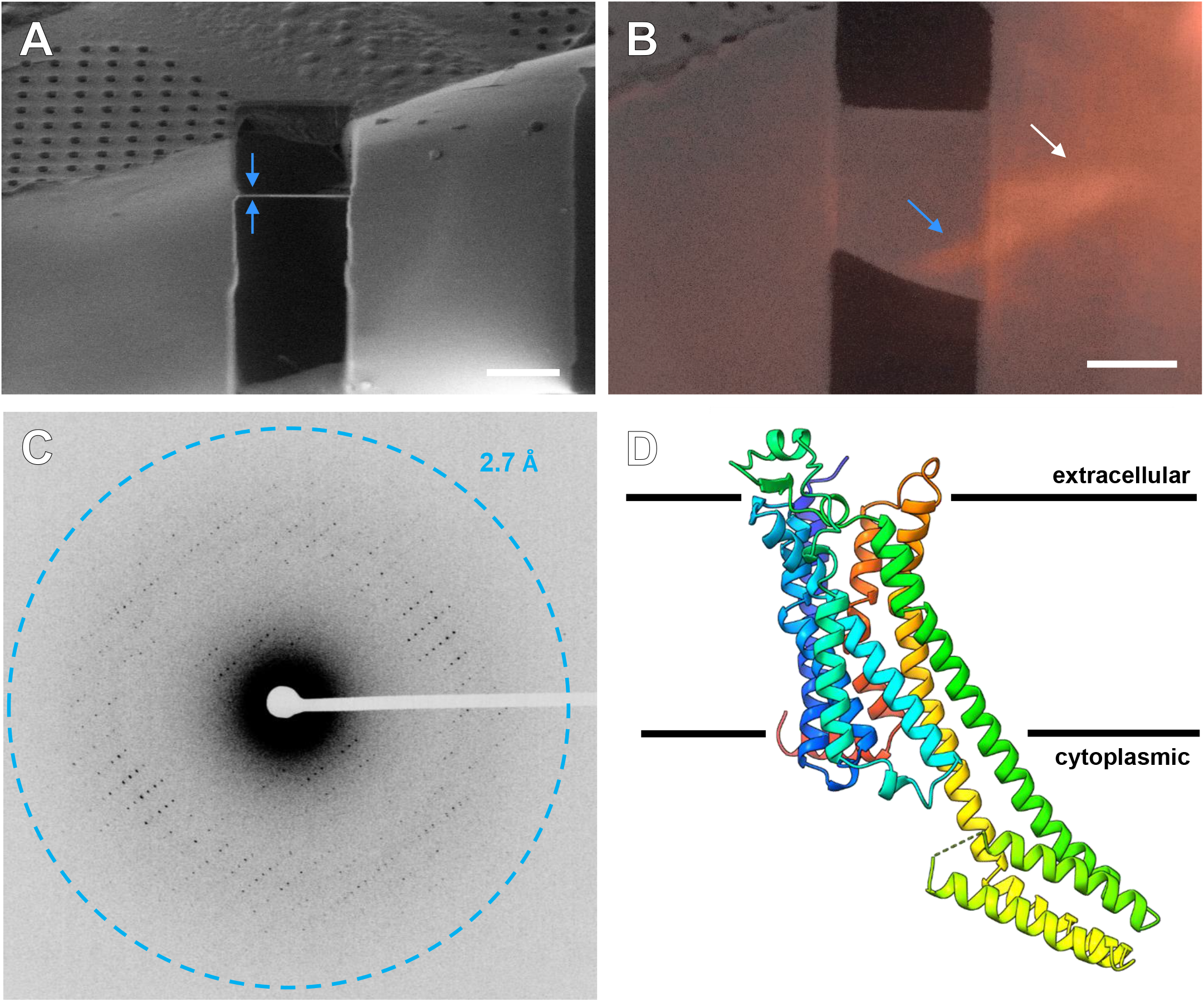
MicroED structure of A_2A_AR from plasma ion-beam milled crystals in thick LCP. (A) Final milled lamella of the GPCR crystal in LCP viewed in the pFIB indicated by blue arrows. The final thickness is approximately 250 nm. (B) Overlaid SEM and iFLM 525 nm fluorescent images of the final lamella confirming the crystal survived the milling process. The blue arrow depicts a milled portion of the crystal, and the white arrow shows an unmilled area of the crystal identified by the fuzzier boundary. (C) MicroED data from the lamella. (D) The 2.7 Å MicroED structure of A_2A_AR determined from a fluorescently labelled, buried microcrystal. Scale bars 10 μm.

The grid containing the A_2A_AR GPCR lamella was transferred into a cryogenically cooled TEM. This lamella was located using low magnification imaging and brought to the eucentric position. A single sweep of continuous rotation MicroED data was collected from a real space wedge between −40° and +40° (**Figure 5C, Methods**). The space group was determined to be C 2 2 21 with a unit cell of (a, b, c) (Å) = (39.04, 177.51, 137.90) and (α, β, λ) (°) = (90, 90, 90) (**Table 2**). The structure was determined by molecular replacement and subsequently refined using isotropic B-factors and electron scattering factors (**Methods**). We observed difference density in the binding pocket corresponding the bound ligand, ZMA. The overall architecture of the protein was as suspected with seven transmembrane helices with a BRIL fusion region in the intracellular region. We did not observe a sodium binding site in the deep pore of our structure (Liu *et al*., 2012), though we cannot rule out its disappearance from either the modest resolution or damage from the electron beam. This structure extended to a resolution of 2.7 Å that is slightly better than our previous results that required changing the phase of the LCP (Martynowycz, Shiriaeva *et al*., 2021), and represents a clear path forward for the routine determination of GPCR crystal structures by MicroED.

**Table 2.**
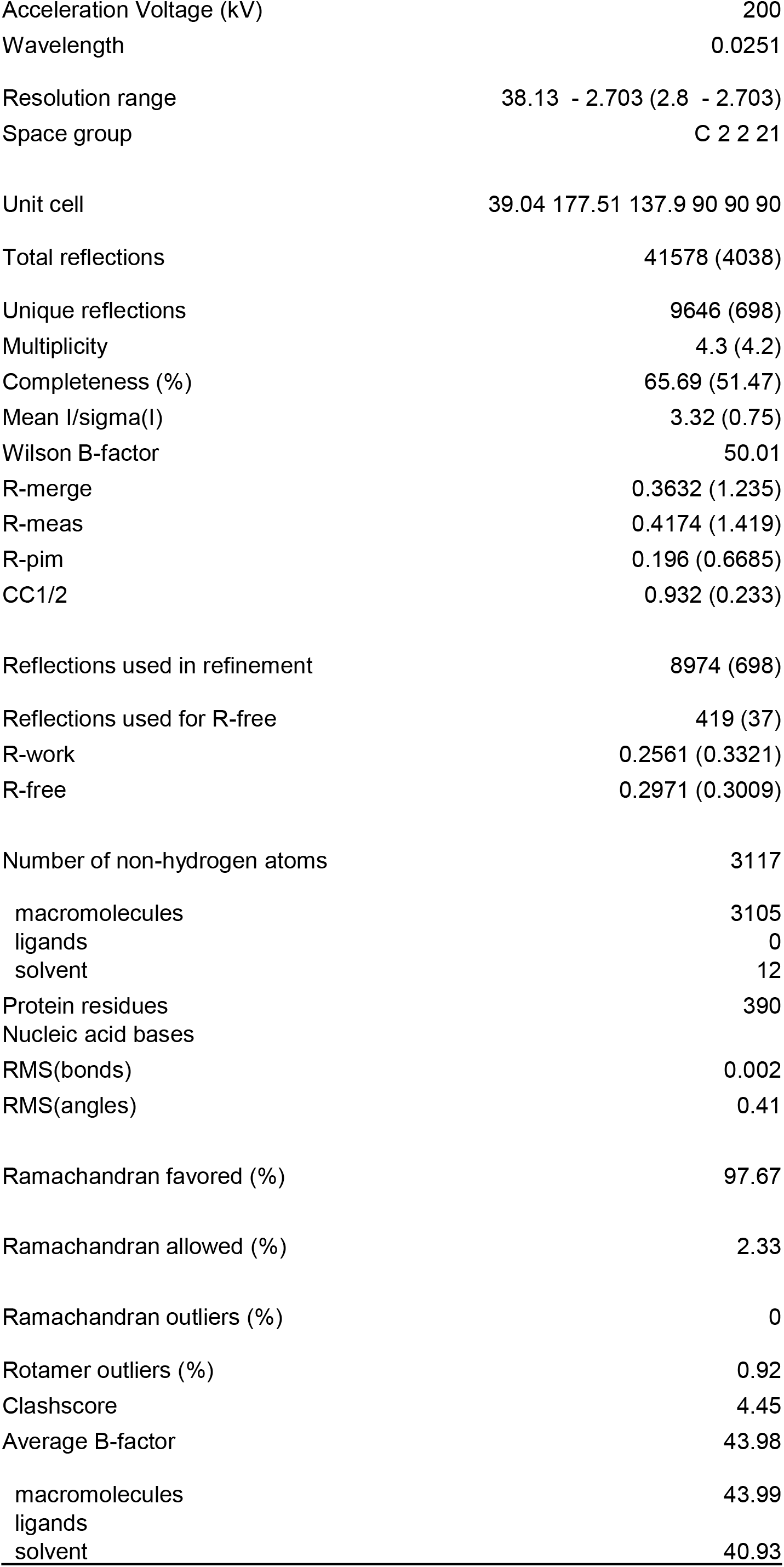
MicroED structure of A_2A_AR determined from a microcrystal buried in LCP

|  |  |
| --- | --- |
| Acceleration Voltage (kV) | 200 |
| Wavelength | 0.0251 |
| Resolution range | 38.13 - 2.703 (2.8 - 2.703) |
| Space group | C 2 2 21 |
| Unit cell | 39.04 177.51 137.9 90 90 90 |
| Total reflections | 41578 (4038) |
| Unique reflections | 9646 (698) |
| Multiplicity | 4.3 (4.2) |
| Completeness (%) | 65.69 (51.47) |
| Mean I/sigma(I) | 3.32 (0.75) |
| Wilson B-factor | 50.01 |
| R-merge | 0.3632 (1.235) |
| R-meas | 0.4174 (1.419) |
| R-pim | 0.196 (0.6685) |
| CC1/2 | 0.932 (0.233) |
| Reflections used in refinement | 8974 (698) |
| Reflections used for R-free | 419 (37) |
| R-work | 0.2561 (0.3321) |
| R-free | 0.2971 (0.3009) |
| Number of non-hydrogen atoms | 3117 |
| macromolecules | 3105 |
| ligands | 0 |
| solvent | 12 |
| Protein residues | 390 |
| Nucleic acid bases |  |
| RMS(bonds) | 0.002 |
| RMS(angles) | 0.41 |
| Ramachandran favored (%) | 97.67 |
| Ramachandran allowed (%) | 2.33 |
| Ramachandran outliers (%) | 0 |
| Rotamer outliers (%) | 0.92 |
| Clashscore | 4.45 |
| Average B-factor | 43.98 |
| macromolecules | 43.99 |
| ligands |  |
| solvent | 40.93 |

### Outlook and Discussion

We present a robust method to determine GPCR protein crystal structures by MicroED from unconverted LCP by targeting buried crystals using cryo-FLM, CLEM, and plasma ion-beam milling. Using this approach, we determined the structure of the human adenosine receptor, A_2A_AR, by MicroED. The protein was fluorescently labelled. Crystals were grown in LCP, looped and then smeared across an EM grid before freezing. The GPCR crystals were buried in dense LCP and could not possibly be identified by using FIB/SEM imaging. Instead, the crystals were located using an integrated fluorescent light microscope. Deep milling through LCP to the depth of the fluorescent crystals was accomplished using a plasma focused ion-beam rather than the traditional gallium beam. To achieve this result, we applied plasma focused ion-beam milling to thin cryogenically frozen biological material. To our knowledge, these are the first biological lamellae milled using plasma focused ion-beam sources for cryoEM experiments in a TEM. The speed of lamellae preparation indicates that xenon mills the fastest with the others following in the order of argon, oxygen, and nitrogen. Although argon milled lamellae faster for the small proteinase crystals, this was because a higher current could be used without destroying these tiny crystals or visibly damaging them during the rough milling steps. For the A_2A_AR crystals buried deep in LCP, the xenon beam at much higher currents could be used, since the possibility of tearing the entire sample away was alleviated. The qualitative metric of lamella cracking suggests that the highest rate of unbroken lamellae occurred using argon or xenon, whereas the lamellae that displayed the least curtaining would by either argon or oxygen. Crystallographic statistics show that the best data is obtained from either argon or xenon, with oxygen and nitrogen performing more poorly. Nitrogen milled lamellae were clear outliers as the worst of all categories overall. Although the oxygen milled lamellae showed better resolution and statistics, the structures from nitrogen or oxygen milled lamellae were of similar overall quality. The MicroED data collected from both argon and xenon milled lamellae of proteinase K were individually of better quality than any data previously recorded from gallium milled lamellae, indicating that there appears to be a clear improvement in data from lamellae prepared by these sources. The improved data quality may arise from reduced damage to the lamellae faces compared to gallium milled lamellae. An improved vacuum also prevents the rapid buildup of amorphous ice in the pFIB/SEM chamber. These improvements are in addition to the increased speed of preparing lamellae using a pFIB, where we manually prepared twenty lamellae across four different sources twice as fast as what could be prepared using a gallium instrument. We suspect that the MicroED data quality could be further improved by polishing the milled lamellae at lower accelerating voltages, as is the standard in materials applications (Stegmann *et al*., 2009). The data collected here represents a first step into the application of plasma beam milling of biological samples for cryoEM investigations. Given the speed and quality of these initial results, we foresee application of this approach to automated lamellae preparation software with throughput gains of up to an order of magnitude over the current state-of-the-art. The improved resolution, data quality, and speed will correspond to improved signal to noise ratios in other cryoEM methods that prepare samples by FIB milling such as cryoET. Furthermore, the approach of plasma ion-beam milling buried membrane protein crystals identified using integrated fluorescence microscopy on the fly will accelerate the adoption of MicroED data collection from critically important membrane proteins.

## Acknowledgements

This study was supported by the National Institutes of Health P41GM136508 and the Department of Defense HDTRA1-21-1-0004. The Gonen laboratory is supported by funds from the Howard Hughes Medical Institute.We would like to thank Abhay Kotecha and Ron Kelley from Thermo Fisher Scientific for their useful discussions and help setting up the pFIB instrument.

## Materials and Methods

### Materials

Proteinase K was purchased from Sigma and used without further purification. Milli-Q water was used for all stock solutions. Cy3 fluorescent dye was purchased from ThermoFisher and used without further purification. All stock solutions were membrane filtered three times. Tetraspecs fluorescent beads were purchased from Invitrogen.

### Protein purification

Expression and purification of A_2A_AR, containing BRIL fusion protein in the third intracellular loop and a C-terminal truncation of residues 317 to 412 (A_2A_AR-BRIL-ΔC), were done as previously described (Jaakola *et al*., 2008; Liu *et al*., 2012).

### Growing protein microcrystals

Proteinase K was crystallized as described (Masuda *et al*., 2017). Protein powder was dissolved at a concentration of 40 mg / mL in 20 mM MES–NaOH pH 6.5. Crystals were formed by mixing a 1:1 ratio of protein solution and a precipitant solution composed of 0.5 M NaNO3, 0.1 M CaCl2, 0.1 M MES-NaOH pH 6.5 in a cold room at 4 °C. Microcrystals grew overnight.

The A_2A_AR, protein was labelled on column with Cy3-NHS ester in accordance with the FRAP-LCP protocol (Fenalti *et al*., 2015). Labelling buffer contained 50mM Hepes pH 7.2, 800mM NaCl, 10% glycerol, 0.025% DDM 0.0025% CHS, 0.1 v/v% Cy3-NHS solution (4 mg/ml in DMF), 100 μM ZM241385. Labelling was carried out for 3 hr at 4 °C. The excess of dye was washed off with the buffer without Cy3-NHS ester. The sample was eluted in 3 cv of elution buffer (50 mM HEPES pH 7.5, 150 mM NaCl, 250 mM imidazole pH 7.5, 0.025 % / 0.005 % (w/v) DDM/CHS, 10% glycerol, 100 μM ZM241385). The complex was concentrated to 30mg/ml with Amicon centrifugal filter units with 100 kDa molecular weight cutoff (Sigma Millipore).

A2AAR-BRIL-ΔC in complex with ZM241385 were reconstituted into LCP by mixing with molten lipid (monoolein:cholesterol 9:1) using a syringe mixer in ratio 2:3.

Crystals for MicroED data collection were obtained in 96-well glass sandwich plates (Marienfeld). Precipitant solution contained 50 to 75 mM sodium thiocyanate, 100 mM sodium citrate pH 4.8, 28% (vol/vol) PEG 400, and 2% (vol/vol) 2,5-hexanediol. Crystals appeared in within 24 hrs and reached full size within 7 days.

#### Grid Preparation

##### Proteinase K grids

Quantifoil Cu 200 R 2/2 holey carbon TEM grids were glow-discharged for 30 s at 15 mA on the negative setting immediately before use. These grids were loaded into a Leica GP2 vitrification robot. The robot sample chamber was loaded with filter paper and set to 4 °C and 95 % humidity for 1 hr before use. 3 μL of protein crystals from the center of the proteinase K tubes were applied to the carbon side of the glow-discharged grid and allowed to incubate for 10 s. Grids were then gently blotted from the back for 10 s. These grids were then immediately plunged into super-cooled liquid ethane. Grids were stored in liquid nitrogen until use.

##### Depth calibration grids

4 μm Tetraspecs fluorescents beads (Invitrogen # T7284C) were diluted to a ratio of 1:10 in a 50% glycerol aqueous solution. 3 uL were pipetted on Quantifoil grids Cu R2/2 (EMS # LFH2100CR2) that were then manually backblotted for 2 seconds and plunge frozen in liquid nitrogen. Alternatively, 0.3 - 0.5 μL droplets were deposited on the grids and the grids were frozen without prior back blotting in order to create domes of glycerol with the Tetraspecs incased in them.

##### A_2A_AR grids

Crystals were looped from glass sandwich plates using a 100 μm MiTeGen dual thickness micromount and carefully transferred to glow-discarged Cu200 R2/2 grids that had been pre-clipped. Looping was conducted under a light microscope next to a humidifier to prevent the LCP from drying out and changing phase during the transfer. Loops full of LCP and crystals were gently slid across the surface of the grid, and the grids were then immediately plunged into liquid nitrogen. All grids were stored at liquid nitrogen temperature before further experiments.

#### Calibrating the milling depth between iFLM and pFIB images

We first used the in-chamber fluorescence light microscopy (iFLM) to localize a thick area containing numerous Tetraspecs at various depths. We milled off the top of the glycerol pile to create a small surface visible in light microscopy, defining our “zero-depth” reference. We registered the depth of the Tetraspecs relative to our zero -depth reference. Then, alternatively milling at high currents by increments of 10 to 4 μm deep and monitoring the disappearance of the Tetraspecs by iFLM allowed us to track the milling depth at which each Tetraspec disappears and compared it to the iFLM-measured depth.

To estimate the depth and then monitor the disappearance of the fluorescent Tetraspecs in the glycerol, fluorescent stacks and reflection stacks were acquired using the iFLM setup in our Hydra: a fixed 20× objective, 0.7 numerical aperture, and working distance of 0.6 mm.

Fluorescence was used to track the beads and reflection was used to monitor the topology of the milling area, mainly to accurately determine the surface of the milled area.

The light source is a 4 LED system (385, 470, 565 and 625 nm) (Thorlabs LED4D242). For fluorescence imaging a fluorescent quad-band filter cube (Semrock LED-DA/FI/TR/Cy5-B-000) is introduced on the light path. For reflective imaging, an empty filter cube is introduced in the light path. The detector is a 3088 x 2064 frame, with a physical pixel size of 2.4 μm. With the ×20 objective, the pixel size is 120 nm.

Stacks of the Tetraspecs were systematically acquire using the fluorescent (excitation wavelength of 470nm) and reflection modalities with the same parameters and a shuttle inclination of 25° resulting in an image normal to the plane of the EM grid. Data was recorded in bin2, with 100% intensity excitation and 1 or 5 ms exposure for each optical slice for the fluorescence and reflection mode, respectively. The Z-step was consistently set to 2 μm.

To mill through the thick glycerol piles, milling was performed with the xenon beam at 30 kV - 15 and 60 nA, at a shuttle inclination of 0° (resulting in a grazing milling angle of 11°) in order to mimic real milling conditions. Milling box X and Z dimensions depended on the size of the glycerol pile. The Y-dimension being the milling step and ranged from 20 to 4 μm (when getting closer to the bead positions). SEM and FIB imaging were done at 500V – 25 pA and 30 kV – 3 pA, respectively. Both used the Everhart-Thornley detector (ETD).

Fluorescence and reflection stacks were combined into multichannel stacks using Fiji (Schindelin *et al*., 2012). To overlay the two images, a Maximal Intensity Projection (MIP) was performed to see all the Tetraspecs present in the stack at once. The first stack acquired was defined as our zero-depth reference and used to estimate the depth of the Tetraspecs encased in the glycerol. The subsequent stacks generated in between each milling increment are then processed the same way and compared to the previous stack. When a Tetraspec disappears between two iFLM stacks associated to a milled depth, the disappearance depth considered was defined as:

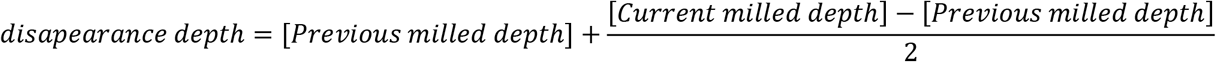

Exceptions were made when it appeared that some of the Tetraspecs were half milled, indicating that the current milled depth is spot on the bead, and therefore the current milled depth was used as the disappearance depth.

Plotting of the Tetraspec depth iFLM estimation vs disappearance depth was done using R and R studio and the following packages: *tidyverse andhere*. As a reference, we included the theoretical FIB-view depth, which is a function of the milling angle relative to the grid plane. In a perfect system where iFLM and milling depth measurements are accurate, the Tetraspecs should disappear at the projected FIB-view depth, defined as:

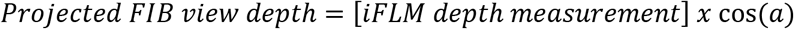

Where a is the milling angle, 11° in this work.

#### Protecting vitrified/biological samples from the plasma beam

Vitrified biological samples need to be protected from the ion-beam during imaging and milling. Inside the FIB/SEM, the sample is typically coated by either a thin layer of pure platinum grains using a sputter coater, a volatile hydrocarbon platinum mixture (Wnuk *et al*., 2009) using a gas injection system (GIS), or a combination of both (Marko *et al*., 2007; Schaffer *et al*., 2017; Martynowycz *et al*., 2019*b*). The sputter-coated platinum layers serve to make the sample conductive and reduce charging artifacts, whereas the thicker GIS deposited platinum protects the sample from the ion-beam during imaging and milling. During these experiments, images are taken using the ion-beam at the lowest current to monitor the sample thickness and adjust for drift or sample movement.

Even at low flux, the ion-beam can damage the sample during imaging (Zhou *et al*., 2019). Milling is typically conducted at much higher beam currents than imaging (Schaffer *et al*., 2017; Beale *et al*., 2020; Martynowycz & Gonen, 2021). For this purpose, it has become a routine to protect the samples by coating the specimens in a thick layer of platinum using a GIS. For many samples, such as mammalian cells that are relatively flat, this approach gives a reasonably smooth, protective layer. For samples with more challenging aspect ratios, such as crystals, the GIS deposition coats differentially because of the facets of the crystals shadowing the grid differently. GIS deposition on grids with crystals often leads to platinum layers that are not homogenous with many blebs, bubbles, and imperfections.

Prior MicroED investigations of thinned crystals all used a gallium focused ion-beam (Polovinkin *et al*., 2020; Zhou *et al*., 2019; Li *et al*., 2018). For many of these samples, a sputter coating of platinum was sufficient to protect the crystals by merely increasing the sputtering time and thereby thickening the layer (60 s – 180 s, ~10 −100 nm of platinum). This approach worked because crystals were milled using very low currents (maximum of typically 300 pA) and the gallium beam size was small enough to overlap very little with the exposed lamella face. This PFIB instrument is not equipped with a sputter coater. Therefore, a new strategy for GIS platinum coating needed to be developed.

At room temperature, the ion or electron beam is used to chemically cleave the volatile mixture, applying material only to the imaged area of the sample (Utke *et al*., 2012). Under cryogenic conditions, the GIS platinum sticks to the grid due to the temperature difference between the volatile, carbon-rich platinum and the cryogenic sample. The rate of platinum deposition at cryogenic temperatures by the GIS is typically too fast to be modified by adding exposure from either the electron or ion beams. After multiple trials, we were able to generate a consistent, dense platinum layer fully protecting all of the crystals along the milling direction. This was accomplished by moving the sample further from the GIS needle to slow the deposition rate and simultaneously imaging the whole grid with a low accelerating voltage, high-current xenon beam (**SI Figure 1**). This coating scheme typically doubled the success rate of lamellae preparation in our hands compared to any conventional method of GIS deposition prior to PFIB milling of these crystals.

Whole grid atlases/montages were created from tiles of individual images taken by the scanning electron microscope operating at an accelerating voltage of either 500V or 2 kV and beam current of 13 pA in the MAPS software (Thermo-Fisher). From the montages, crystals were selected that were not within a few μm of a grid bar, nor within 3 grid squares of the edge of the grid. Dozens of crystals across over ten grids were identified in this way to test various platinum protection setups and testing of milling strategies. After many failures, the results of the beam comparison within this investigation were conducted, where 20 crystals were identified under these criteria.

#### Machining proteinase K microcrystals using the pFIB

The twenty crystals were divided into four equal size groups of five, one group for each plasma ion-beam to test. For each gas source, crystals were milled sequentially using identical milling strategies—the same preset pattern, beam currents, and times. The Z-depth of the patterns and currents were the only parameters adjusted. For each crystal and gas source, the milling was conducted in four steps with approximately the same parameters summarized in Supplementary Table 2. Generally, all milling was conducted using the cleaning cross section for each pattern with 85% overlap between both X and Y spot positions. The first milling step used an approximately 1 nA current to mill two boxes of 6 × 6 um separated in the middle by 2 μm. The second utilized two cleaning cross sections of 5 × 1 μm in size separated by 1μm that used an approximately 0.3 nA current. The third step consisted of two 5 × 0.5 μm boxes separated by 0.5 μm with a beam current of 0.1 nA. The final milling step consisted of two 5 × 0.3 μm boxes separated by 300 nm that were milled with a beam current of approximately 30 pA. Currents between sources were chosen to be within 1 aperture number from the prior source to minimize the number of alignments between experiments (**SI Table 2**). All cleaning cross sections above the lamella were milled from the top to the bottom, whereas all the cross sections below the lamella were milled from the bottom to the top. The sputtering rate for a drawn pattern in the microscope software is set to solid silicon, which is much denser than vitrified water or biological materials. We empirically determined reasonably adapted milling times by varying the dictated Z dimension, or depth of the drawn patterns. For xenon, we used a depth setting of 5, 3, 2, and 2 μm deep for each step. For argon, these were 6, 3, 2, 2 μm deep. For nitrogen, we used 20, 10, 4, 4 μm. Finally, we used 8, 6, 4, 4 μm for the oxygen beam. These settings are summarized in **SI Table 1**. In most cases, these were higher than strictly necessary to ensure second passes would not be needed. However, even with the depths used, the nitrogen lamellae required constant manual intervention that was still unable to rescue some of the lamellae. Typically, argon and xenon lamellae were completed with total milling times of between 4 and 20 mins depending on alignments between milling steps and various manual microscope operations. Each nitrogen lamella took approximately 15 - 30 mins of on-sample milling time. For oxygen, this was similarly 15-30 mins per lamella. A complication to the timing was the manual operation and shortcomings of specific gasses. For example, focusing the argon and xenon beams is more challenging than a gallium beam, but relatively simple. The oxygen and nitrogen beams are very difficult to focus and align at low beam currents. Positioning lamellae was also much easier for the heavier ions since the focused images were much sharper in general. Finally, imaging lamellae using the various ion beams changes the contrast in the electron beam due to the differential breakdown of the GIS deposited platinum over time and differing by each ion. For example, oxygen lamella #2 (**Supplementary Figure 6**) was all but invisible after milling, and even after repeated attempts, the SEM image had to be zoomed out to even understand where the lamella was located. In our experiments, the contrast changing of the GIS deposited platinum without the ion-assisted deposition described herein was much worse, essentially making many attempts at milling with nitrogen or oxygen much more challenging than simply using a gallium beam source.

##### Identification and machining of A_2A_AR crystals

Frozen A_2A_AR grids were transferred into the pFIB/SEM under cryogenic conditions. All-grid montages were collected using the SEM operating at 500 V prior to platinum coating. The low accelerating voltage prevents damaging the sample and allows for visualization of the sample with improved contrast compared to the platinum coated sample. After coating, almost all images in the SEM appear similar. Areas of thick LCP seen in the SEM were inspected in the iFLM using either the 535nm fluorescent signal or the reflected signal. We quickly found that the reflected signal at all wavelengths did not allow us to identify any crystals, whereas the fluorescent signal of Cy3 was simple to identify, with sharp edged crystals visible at various depths of the LCP hills. At each area of interest, a stack of images was obtained in both reflective mode at a single wavelength of 535 nm, and then using all four wavelengths using the fluorescent filter. The range of image steps in the Z direction were defined by first identifying the top of the LCP and the grid surface in reflective mode. This total thickness was then rounded to the nearest micron and imaged at 2 μm Z intervals. The image where the crystal of interest was visually sharpest was taken to be the true depth of the crystal. Images taken in the light microscope were correlated to images taken at the mapping position in the SEM using the MAPS software. The milling locations in the pFIB were created by locating a feature in the correlated Multiple positions were screened until a crystal was identified that was not on top of a grid bar or within the projected range of a grid bar, and would not be occluded by features after milling to the prescribed depth. The milling was conducted using the xenon beam, where the first pattern below the sample with a 2 μm offset was 10 μm wide, 30 μm tall, and 40 μm deep milled at a current of 15 nA. The top pattern was then milled at a 2 μm offset from the crystal and was 10 μm wide, 10 μm tall, and 30 μm deep milled at a current of 15 nA. The rest of the steps were conducted using an x,y,z size of 10, 1, 10 μm currents of 1nA, 0.3nA, and 0.1nA. The final lamellae was monitored in the SEM using a 2kV accelerating voltage and 13pA current with intermediate images were taken between each cut of the pFIB beam. The milling was halted when the contrast of the lamellae flipped in the SEM image.

#### MicroED data collection

Grids containing milled proteinase K crystals were rotated such that the TEM rotation axis was 90° from the plasma-beam milling axis. The grids were then loaded into a cryogenically cooled Thermo-Fisher Titan Krios 3Gi transmission electron microscope operating at an accelerating voltage of 300 kV. Low magnification montages of each grid were collected at a magnification of 64 × and used to locate the milled lamellae. Each lamella was brought to its eucentric position before data collection. MicroED data were collected by continuously rotating the stage at a rate of approximately 0.15° / s for 420 s, covering a total rotation range of approximately 63°, respectively. This typically spanned the real space wedge corresponding to approximately −31.5° to +31.5°. Data were collected using a 50 μm C2 aperture, a spot size of 11, and a beam diameter of 20 um. Under these conditions, the total exposure to each crystal was approximately 1.0 e- Å-2. Diffraction data were collected from a small, isolated area from the middle of each lamella of approximately 2 μm in diameter using the 100 μm selected area aperture to remove unwanted background noise. All data were collected using twofold binning and internally summed such that each image recorded a 0.5 s exposure spanning approximately 0.075° of rotation. In this way, each image stack contained 840 images, the last of which was discarded for having an unequal number of frames. A single sweep of continuous rotation MicroED data was collected from each lamella.

For A_2A_AR, MicroED data was collected on a Talos Arctica operating at liquid nitrogen temperatures at an accelerating voltage of 200 kV. Data were collected by continuously rotating at a rate of 0.5 °/s for 160 s, spanning a real space wedge from −40 ° to +40 °. Data were collected on a CetaD CMOS 4096 x 4096 detector operating in rolling shutter mode with correlated double sampling active.

#### MicroED data processing

Movies in MRC format were converted to SMV format using a parallelized version of the MicroED tools (https://cryoem.ucla.edu/downloads). Each proteinase K dataset was indexed and integrated using XDS in space group 96. The A_2A_AR dataset was indexed in DIALS (Winter *et al*., 2018), and then integrated in XDS. All datasets were scaled using XSCALE. For merging all the proteinase K data, xscale_isocluster was used. Datasets that were of either much poorer resolution or scaling correlation below 90% were discarded. For all crystals, the space group was verified using POINTLESS. Data were merged without scaling using AIMLESS, the subsequent intensities were converted to amplitudes in CTRUNCATE, and a 5% fraction of the reflections were assigned to a free set using FREERFLAG (Winn *et al*., 2011).

In order to achieve the best model possible from our collected data and to test if data derived from different ion sources could reasonably be merged together, we created an additional merged data set from across all the lamellae. First, a naïve merge of all the integrated datasets was conducted. To identify which datasets from each source merged best, isoclustering (Assmann *et al*., 2020) was performed. Poorly contributing data was discarded and the remaining datasets were automatically assigned weights and merging order to yield a“Best Merge” from this subset (**Figure 3, Table 1**). This merged dataset was composed of 12 of the 20 individual datasets – 5 argon datasets, 5 xenon datasets, and 2 oxygen datasets. This final merged dataset had overall statistics superior to any of the individual datasets or subsets of merged data from individual gases and to a slightly better resolution (**Table 1**). The structure of proteinase K was determined from this dataset and refined identically to the other sources. The refined structure from this merged data had an overall R work and R free of 11.92 / 16.34. These statistics were better than the models derived from argon and xenon alone. The results suggest that merging data from across different ion sources is possible without degradation of the model. It could be that there is some benefit in merging data between the sources given the improved metrics, however it is difficult to separate the improvements in statistics and resolution from the increase in redundancy.

#### Structure solution and refinement

The structures of proteinase K were determined by molecular replacement in PHASER using the search model 6cl7. The structure of A_2A_AR was determined by molecular replacement using 4EIY as a search model. The solutions were refined in Phenix.refine. For proteinase K models, the first refinements used isotropic B-factors and automatic water picking that resulted in an Rwork / Rfree of approximately 0.18/0.20. The refined model was inspected in Coot. Several Calcium and NO3 ions were placed in the difference maps, an incorrectly assigned residue (SER^312^ ->ASP^312^) was fixed, and alternative conformations were identified for several residues. Occupancies were refined for nitrate ions and alternate side chain conformations. This model was refined again in Phenix using the same settings that resulted in approximate Rwork/Rfree of 0.16/0.19. After another visual inspection in Coot, the model was refined again in Phenix using automatic water picking and anisotropic B-factor refinement for all atoms that resulted in Rwork/Rfree of 0.15/0.18. From here, the model was refined again in REFMAC5 using automatic matrix weights, anisotropic B-factors, and added hydrogens, where the final Rwork/Rfree dropped to 0.12/0.16. The A_2A_AR model was refined in PHENIX.REFINE using isotropic B-factors to a final Rwork / Rfree of 25/30 and resolution of 2.7 Å.

#### Figure and Table preparation

Figures were prepared using ChimeraX (Goddard *et al*., 2018), FIJI (Schindelin *et al*., 2012), the matplotlib package in Python 3.6 in a Jupyter notebook and R. Figures were arranged in PowerPoint, and Tables were arranged in Excel. Maximum intensity projections were calculated in FIJI (Schindelin *et al*., 2012).

## Supplementary Information

### Supplementary Table and Figure Legends

**Supplemental Figure 1.**
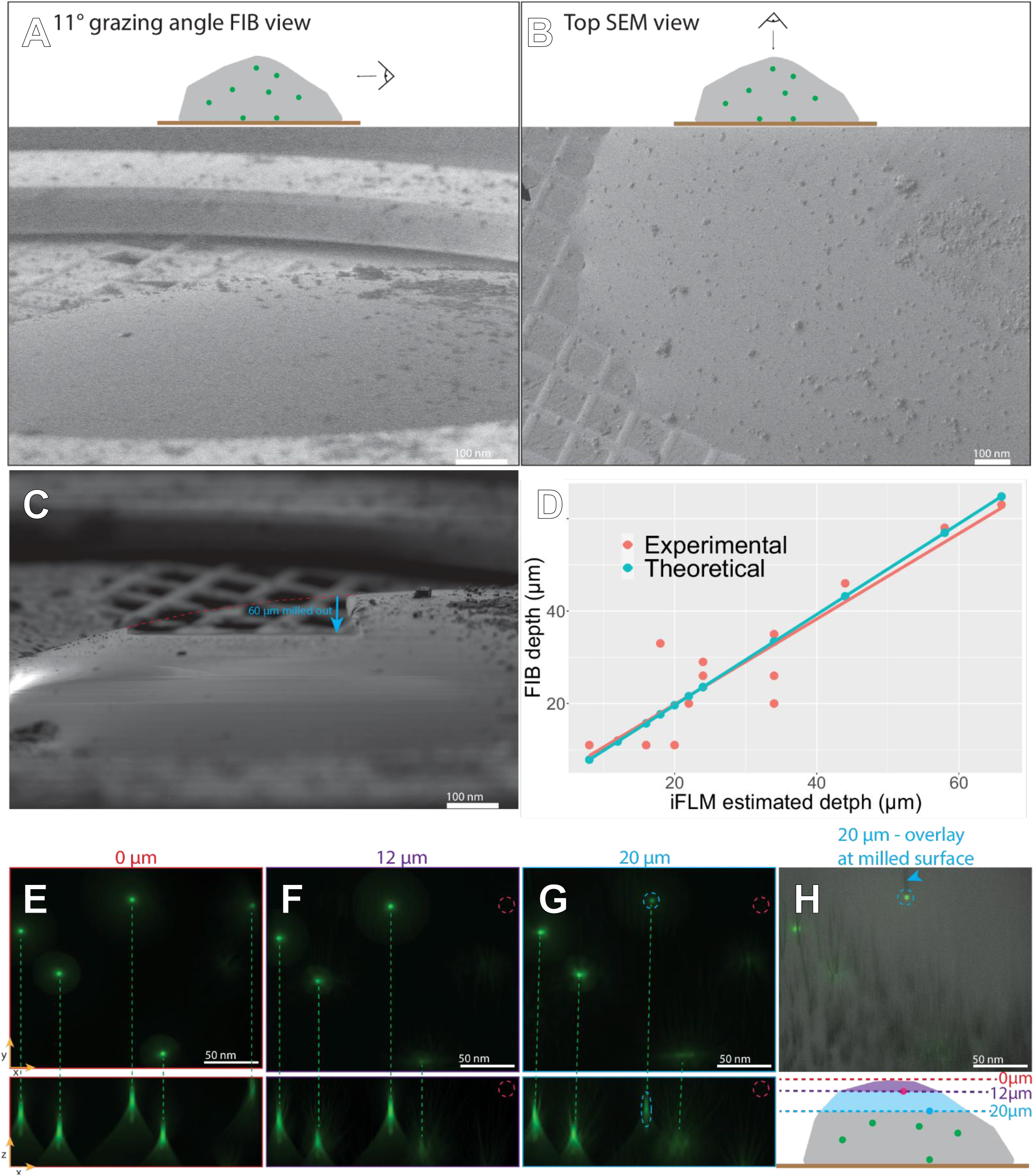
Correlation between the iFLM and pFIB measured depth. (**A**) <u>Top panel</u>: Cartoon of a glycerol pile (gray) laying on a grid (orange) with 4 um Tetraspecs encased in it (green). The eye and arrow represent the grazing angle at which the imaging/milling is done with the FIB gun. Bottom panel: FIB view using Argon at 6pA – 30kV showing the frozen glycerol pile. (**B**) <u>Top panel</u>: Same cartoon as in (A) with the eye and arrow representing the top view acquired from the electron gun. Bottom panel: SEM view at 25pA – 500V of the pile of glycerol. (**C**) FIB view at 4nA – 30kV and same angle as in (A) after multiple incremental milling steps. Here, 60 μm have been milled in total (blue arrow). The red outline shows the initial curve of the glycerol pile. (D) Plot of the iFLM measured depth (x-axis) versus the FIB measured depth (y-axis). (**E, F and G**) <u>Top panel</u>: Maximum intensity projection of an X-Y oriented stack acquired at the milling site. <u>Bottom panel</u>: Maximum intensity projection of an X-Z oriented stack acquired at the milling site. (E) was acquired before any milling was performed. It is the zero reference. (F and G) were acquired after milling 12 μm and 20 μm, respectively. The dashed purple circles show a Tetraspec that disappeared after 12 μm of milling. The yellow dashed purple circles represent a Tetraspec that is spot-on 20 μm deep. (**H**) Overlay optical slice at 20 μm deep showing the surface of the lamella in reflective mode (gray) and the fluorescent Tetraspecs (green). The bead circled in blue sits on the milled surface, its milled “shadow”, creating a curtaining artefact can be seen behind it (blue arrowhead). (**I**) Same cartoon as in (A) and (B) with dashed lines corresponding to the different depths at which the iFLM stacks showcased in (E, F and G) were acquired.

**Supplemental Figure 2.**
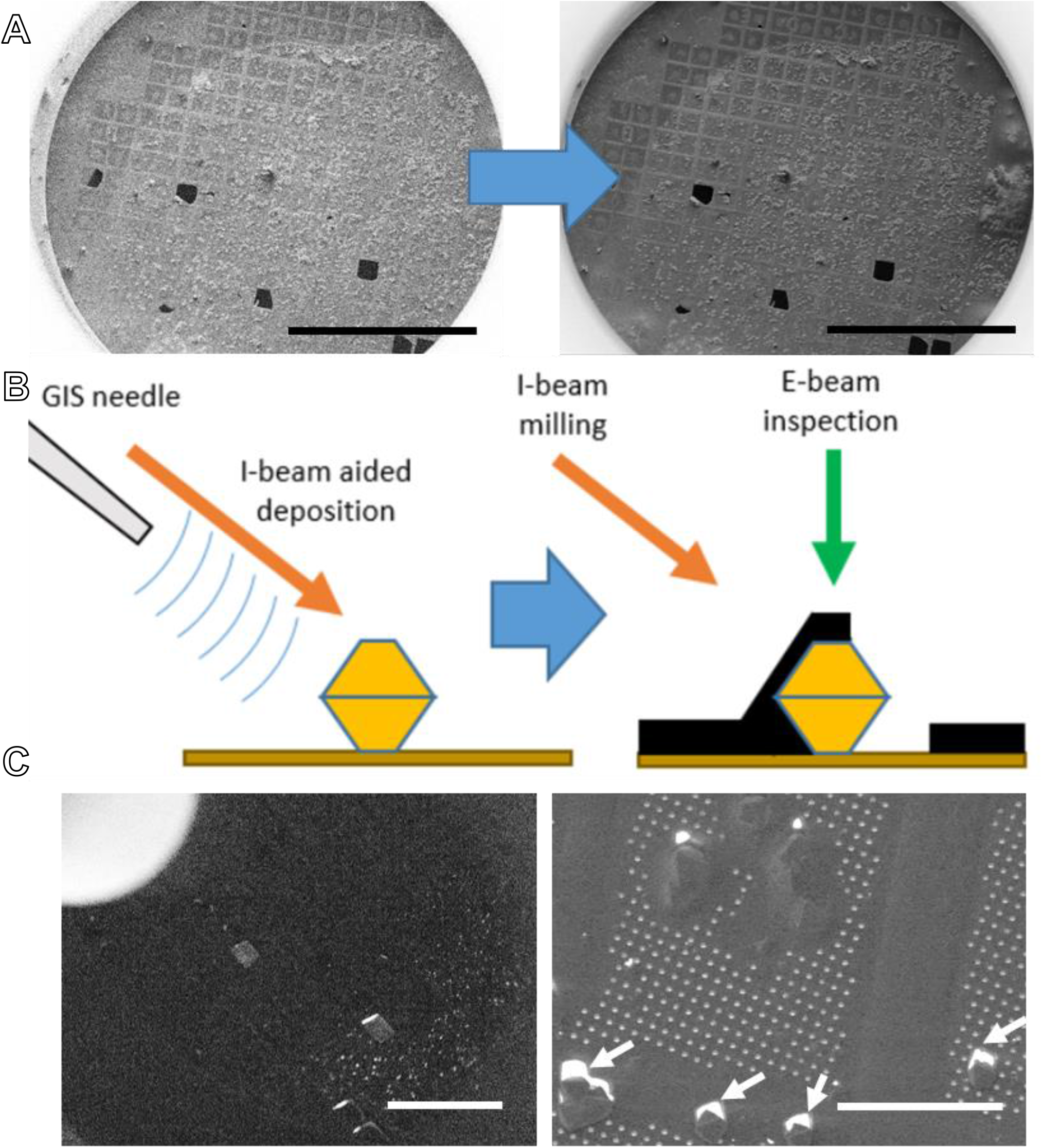
Grazing incidence GIS platinum deposition aided by the plasma ion beam. (A) whole-grid SEM image before (left) and after (right) GIS deposition. (B) Cartoon depiction of ion-assisted GIS platinum deposition (left) and how the geometry leaves the back of the crystal shadowed (right). (C) The left hand image shows the view in the xenon ion-beam during the GIS platinum coating of the grid, and the right hand side shows the grid after coating with clear uncoated areas behind each crystal.

**Supplemental Figure 3.**
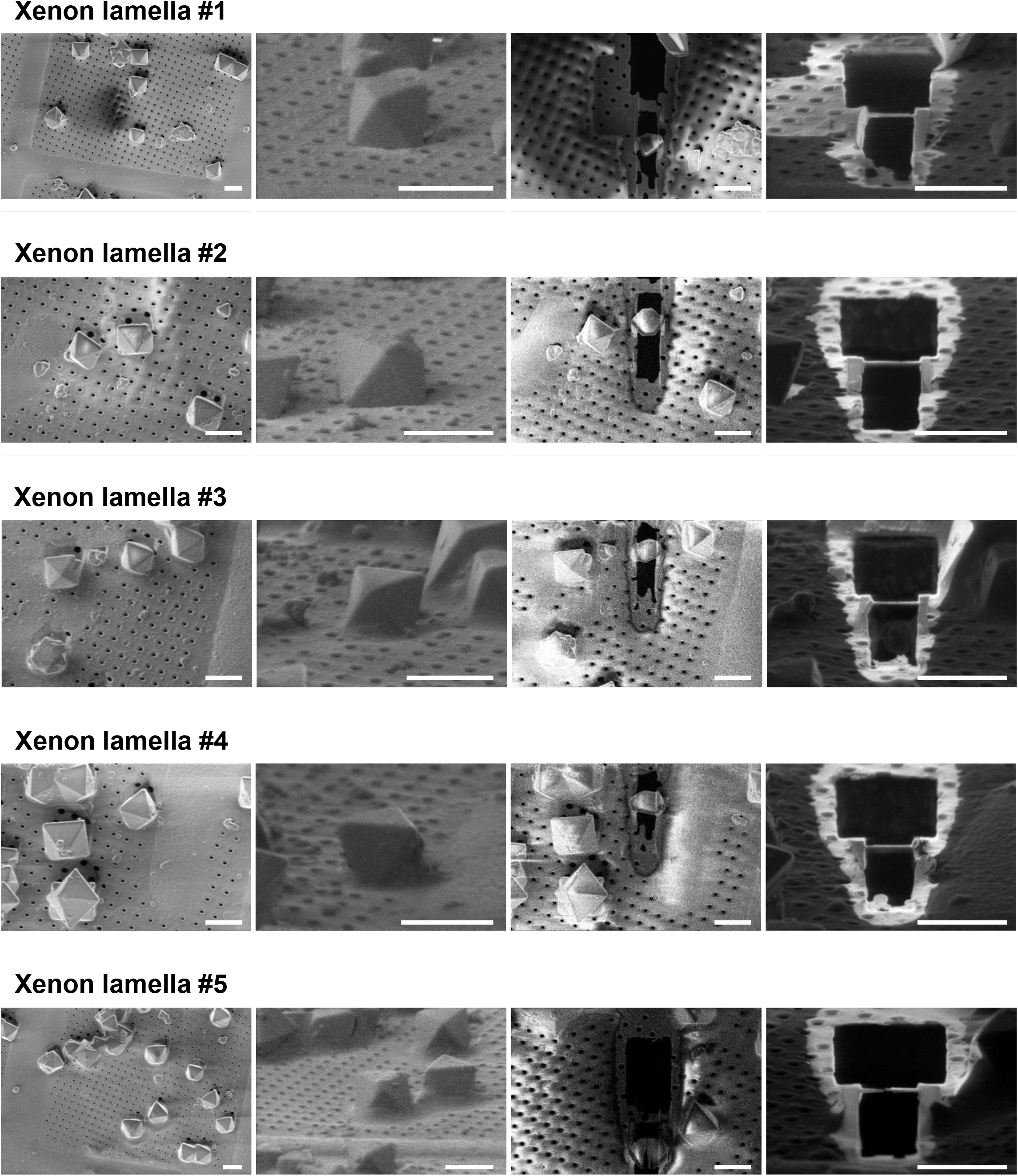
SEM and pFIB images of proteinase crystals milled using the xenon beam. All scale bars are 10 μm.

**Supplemental Figure 4.**
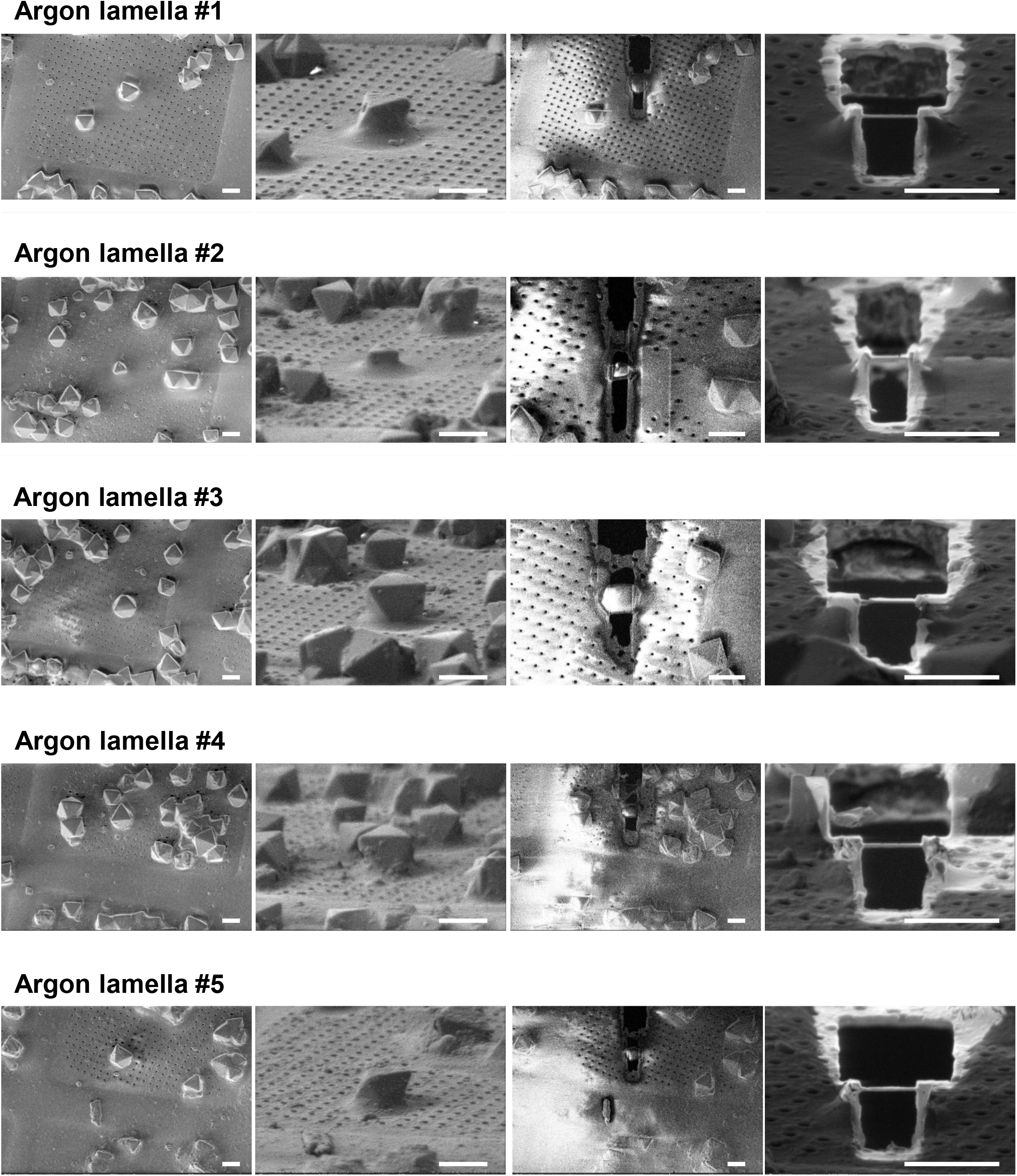
SEM and pFIB images of proteinase crystals milled using the argon beam. All scale bars are 10 μm.

**Supplemental Figure 5.**
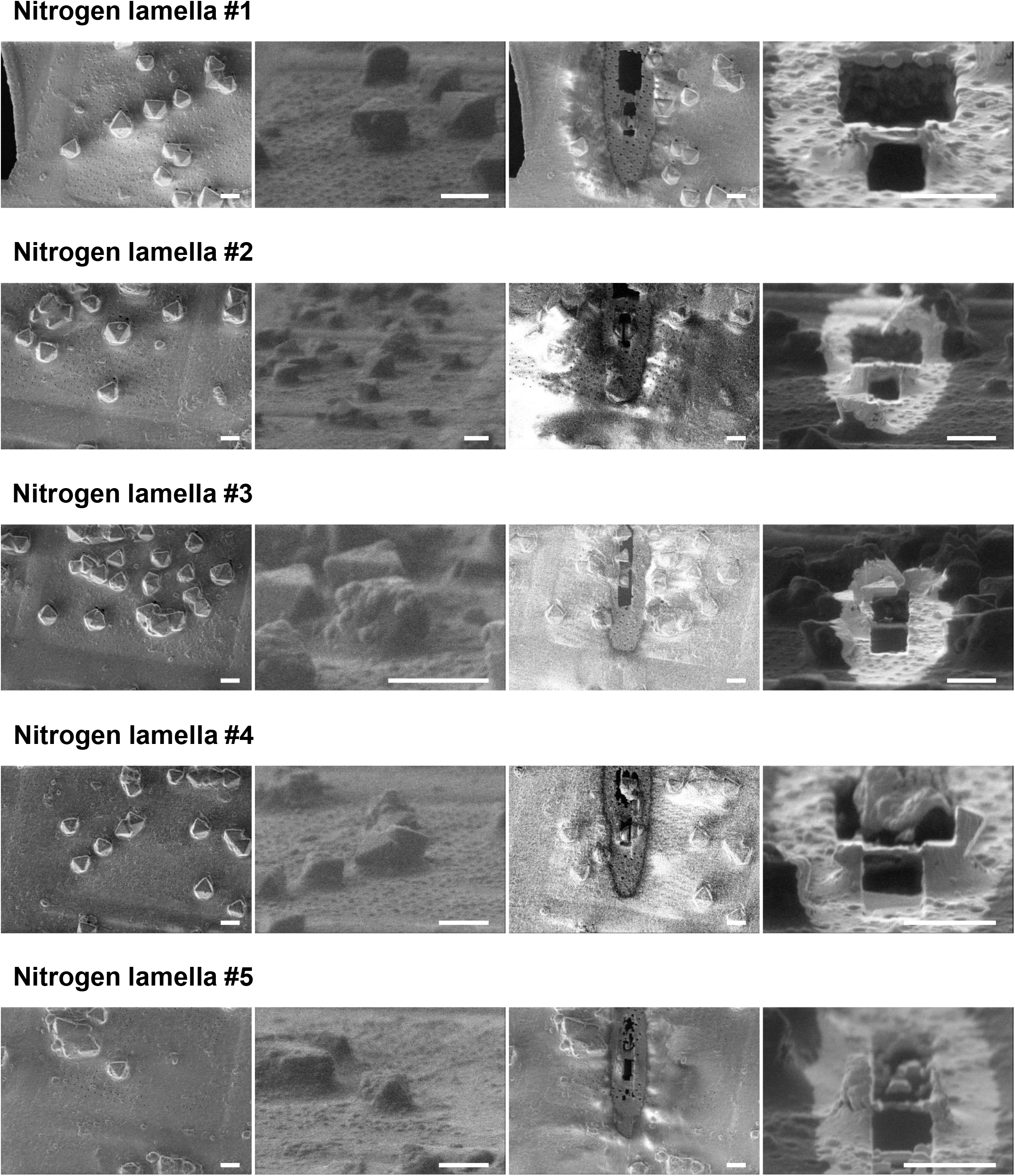
SEM and pFIB images of proteinase crystals milled using the nitrogen beam. All scale bars are 10 μm.

**Supplemental Figure 6.**
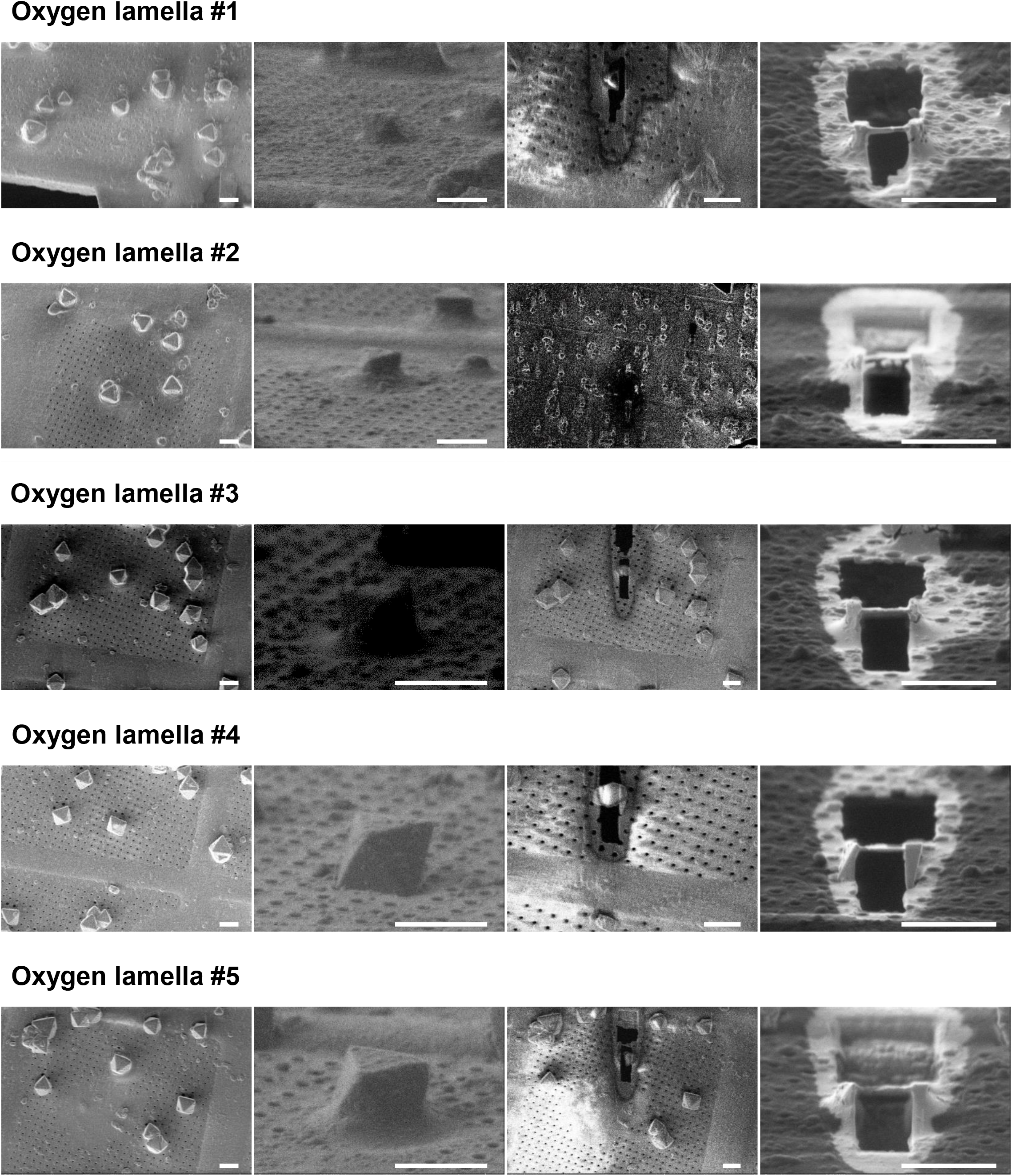
SEM and pFIB images of proteinase crystals milled using the oxygen beam. All scale bars are 10 μm.

**Supplemental Figure 7.**
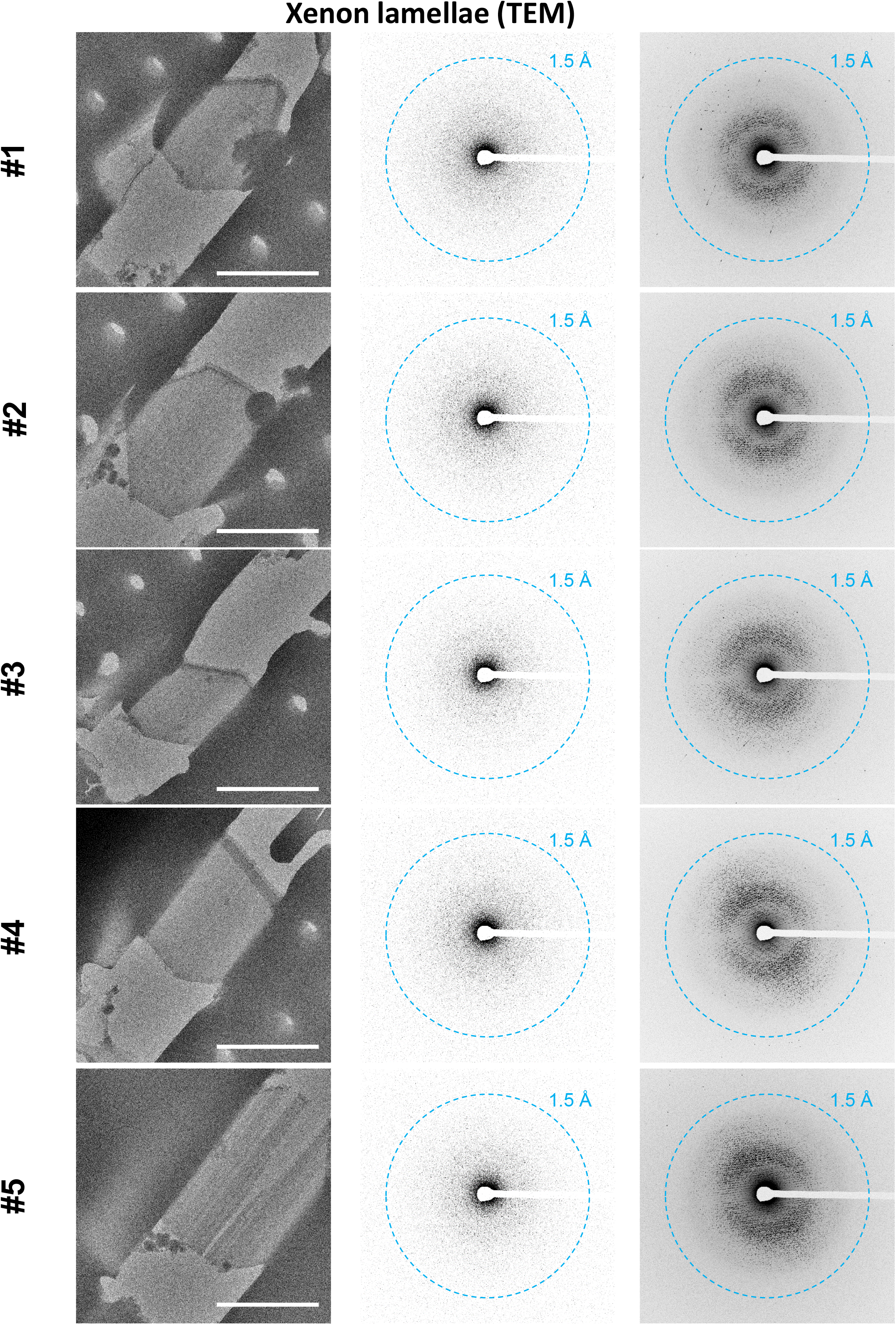
TEM images of proteinase crystals milled using the xenon beam. All scale bars are 10 μm. Single diffraction images from each movie are depicted in the center, and the maximum intensity projections shown on the right.

**Supplemental Figure 8.**
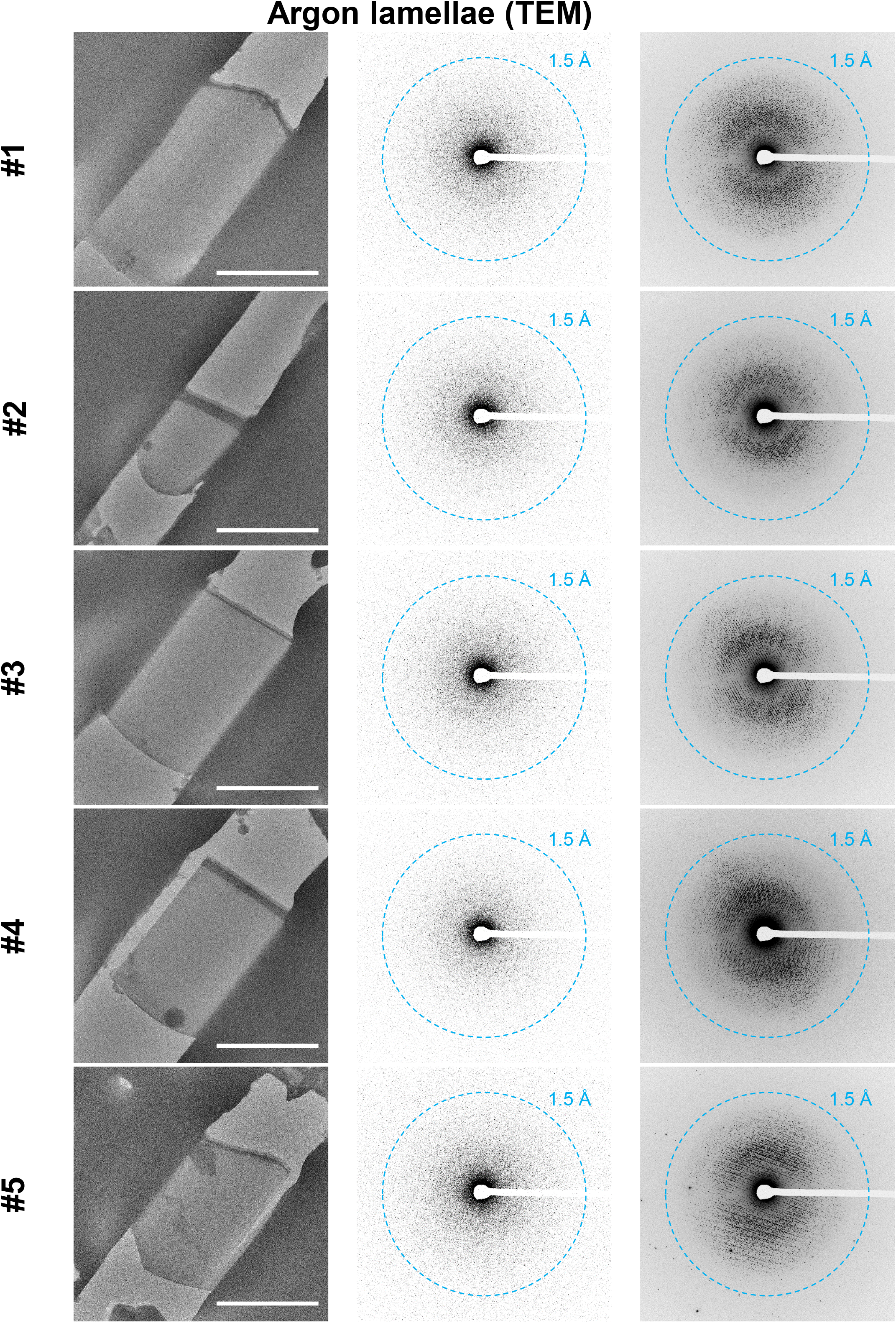
TEM images of proteinase crystals milled using the argon beam. All scale bars are 10 μm. Single diffraction images from each movie are depicted in the center, and the maximum intensity projections shown on the right.

**Supplemental Figure 9.**
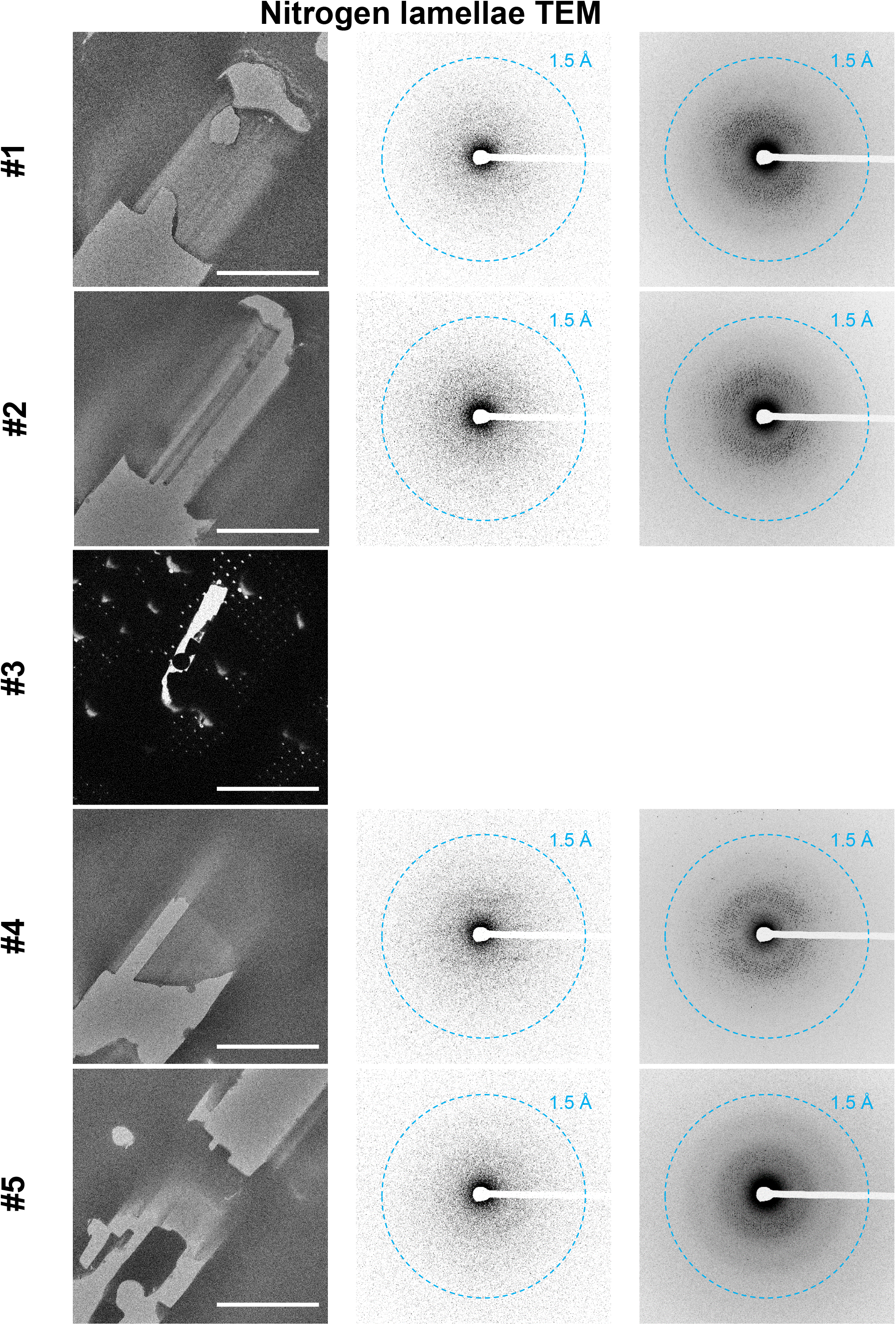
TEM images of proteinase crystals milled using the nitrogen beam. All scale bars are 10 μm. Single diffraction images from each movie are depicted in the center, and the maximum intensity projections shown on the right.

**Supplemental Figure 10.**
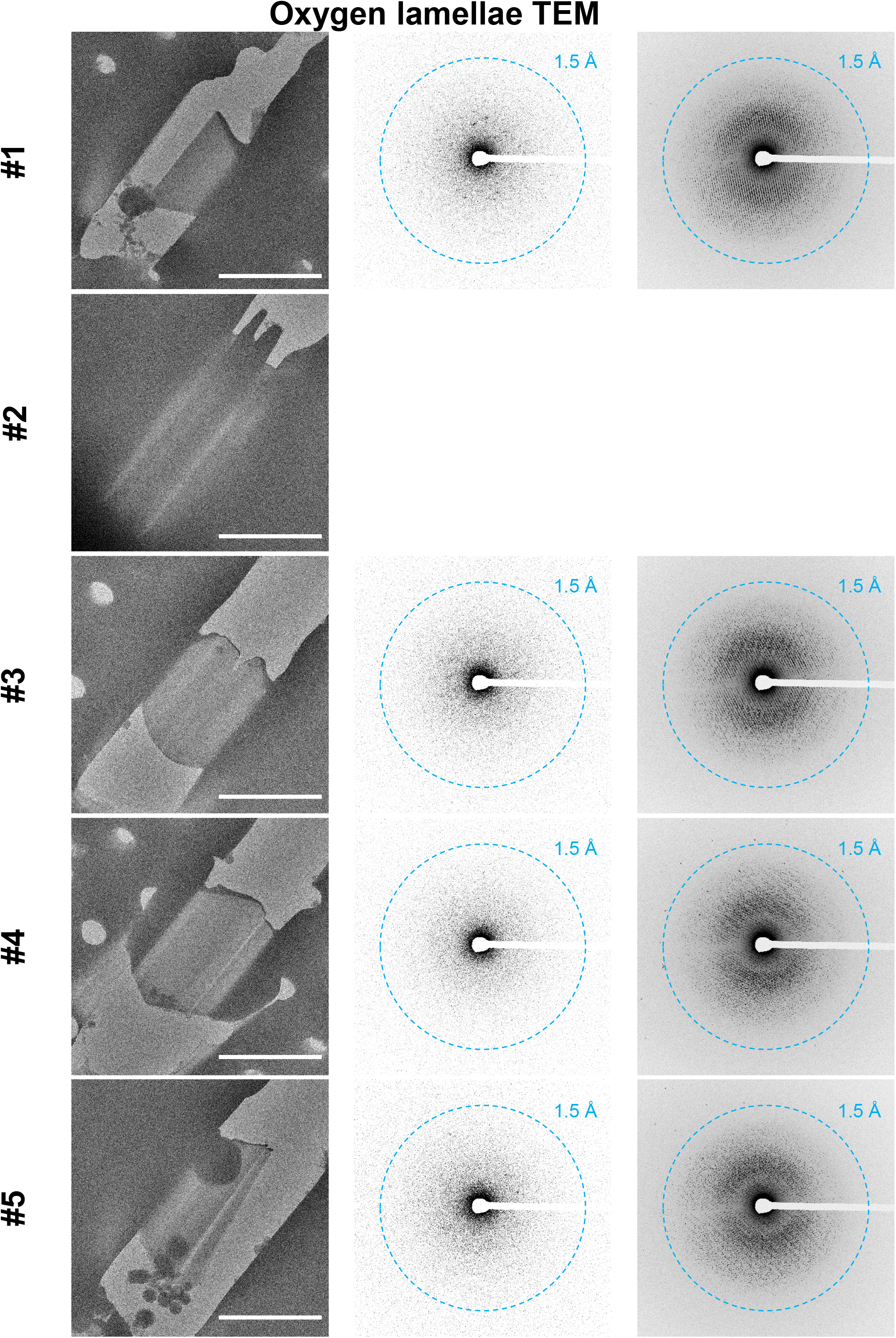
TEM images of proteinase crystals milled using the oxygen beam. All scale bars are 10 μm. Single diffraction images from each movie are depicted in the center, and the maximum intensity projections shown on the right.

**Supplemental Figure 11.**
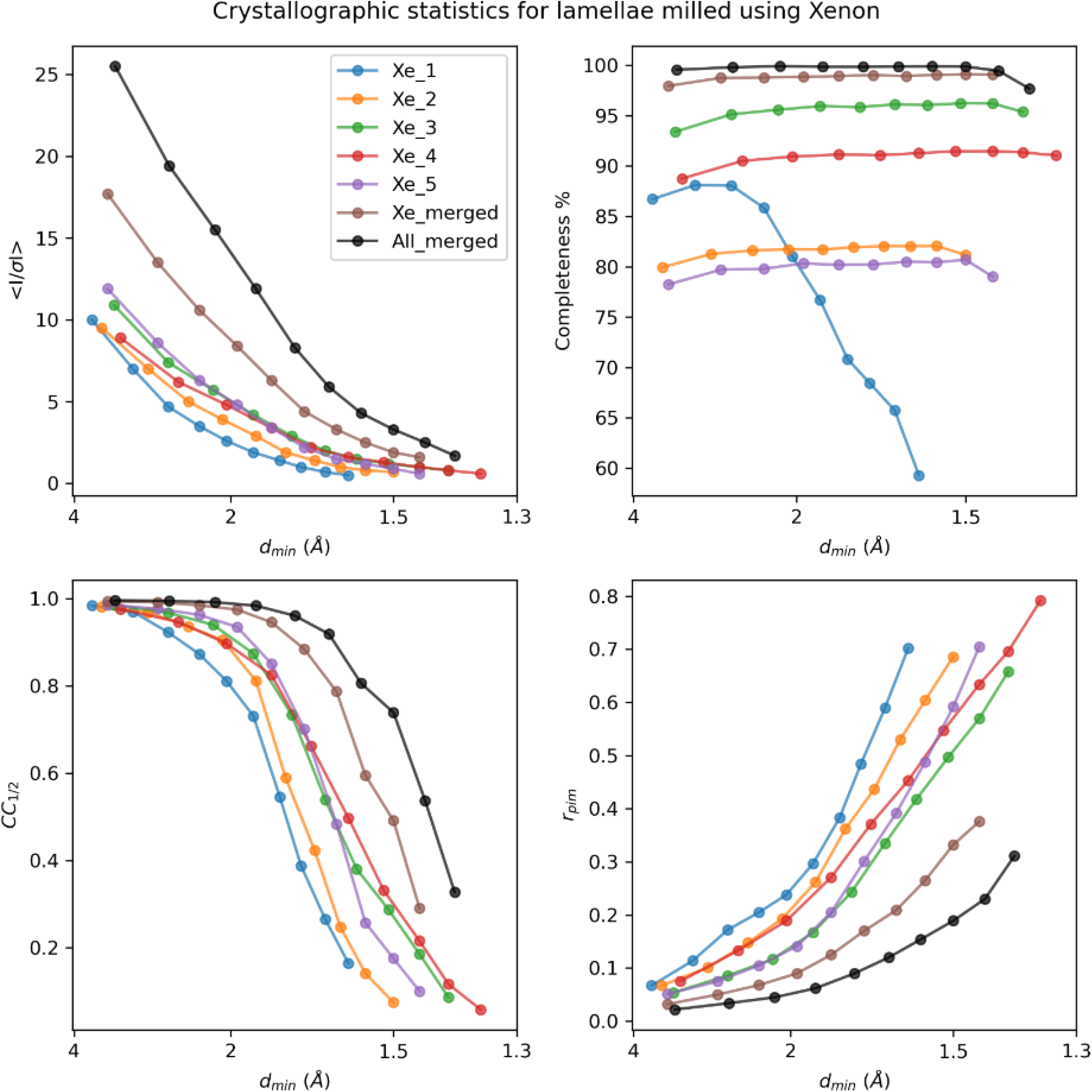
Crystallographic statistics for xenon ion-beam milled lamellae. Plots depict the mean signal to noise ratio (<I / σ (I)>) (top left), completeness (%) (top right), mean half-set correlation coefficient (CC_1/2_) (bottom left), and multiplicity corrected R factor (Rpim)(bottom right) as functions of the dmin resolution bins (Å). The “best merge” data set is included for comparison in each case.

**Supplemental Figure 12.**
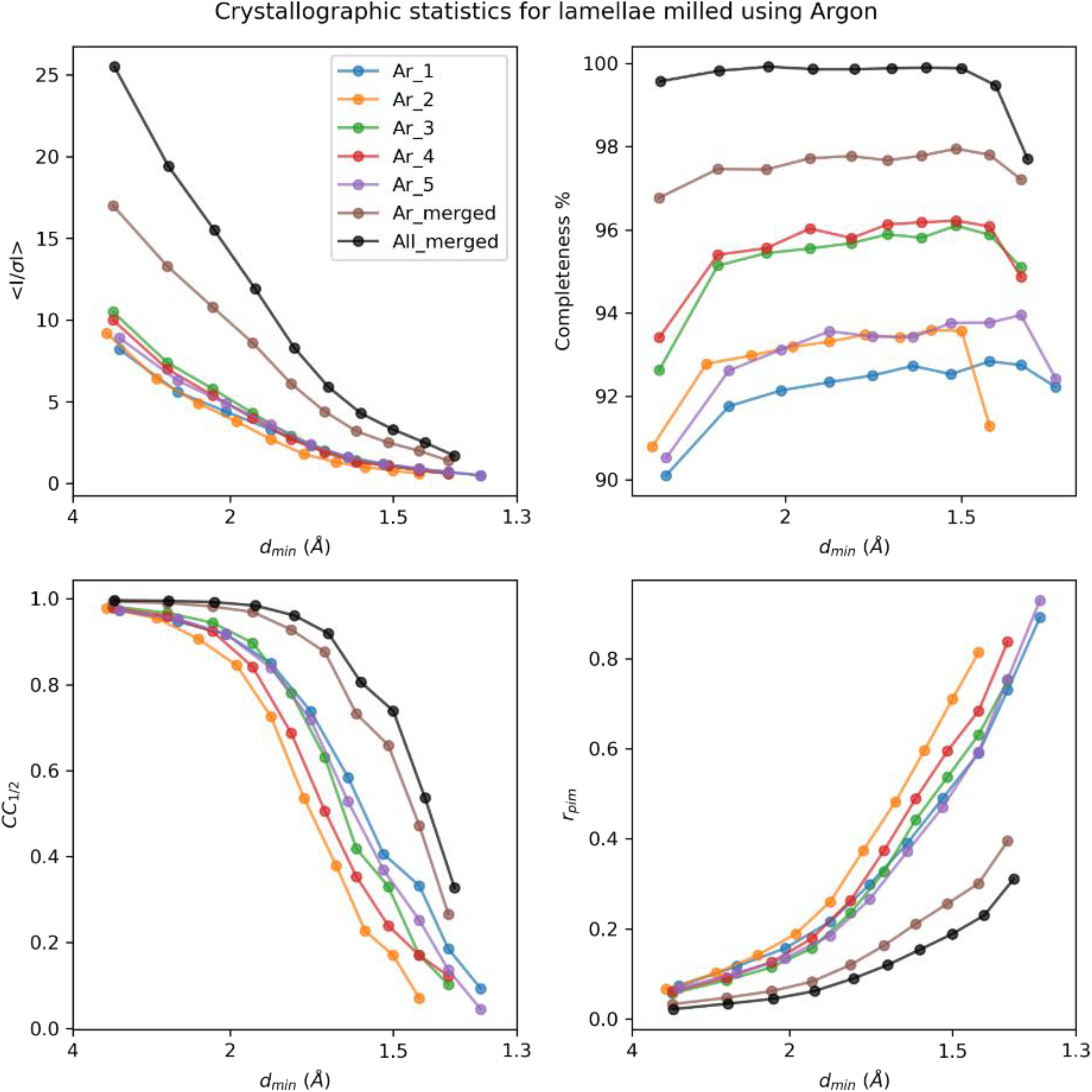
Crystallographic statistics for argon ion-beam milled lamellae. Plots depict the mean signal to noise ratio (<I / σ (I)>) (top left), completeness (%) (top right), mean half-set correlation coefficient (CC_1/2_) (bottom left), and multiplicity corrected R factor (Rpim)(bottom right) as functions of the dmin resolution bins (Å). The “best merge” data set is included for comparison in each case.

**Supplemental Figure 13.**
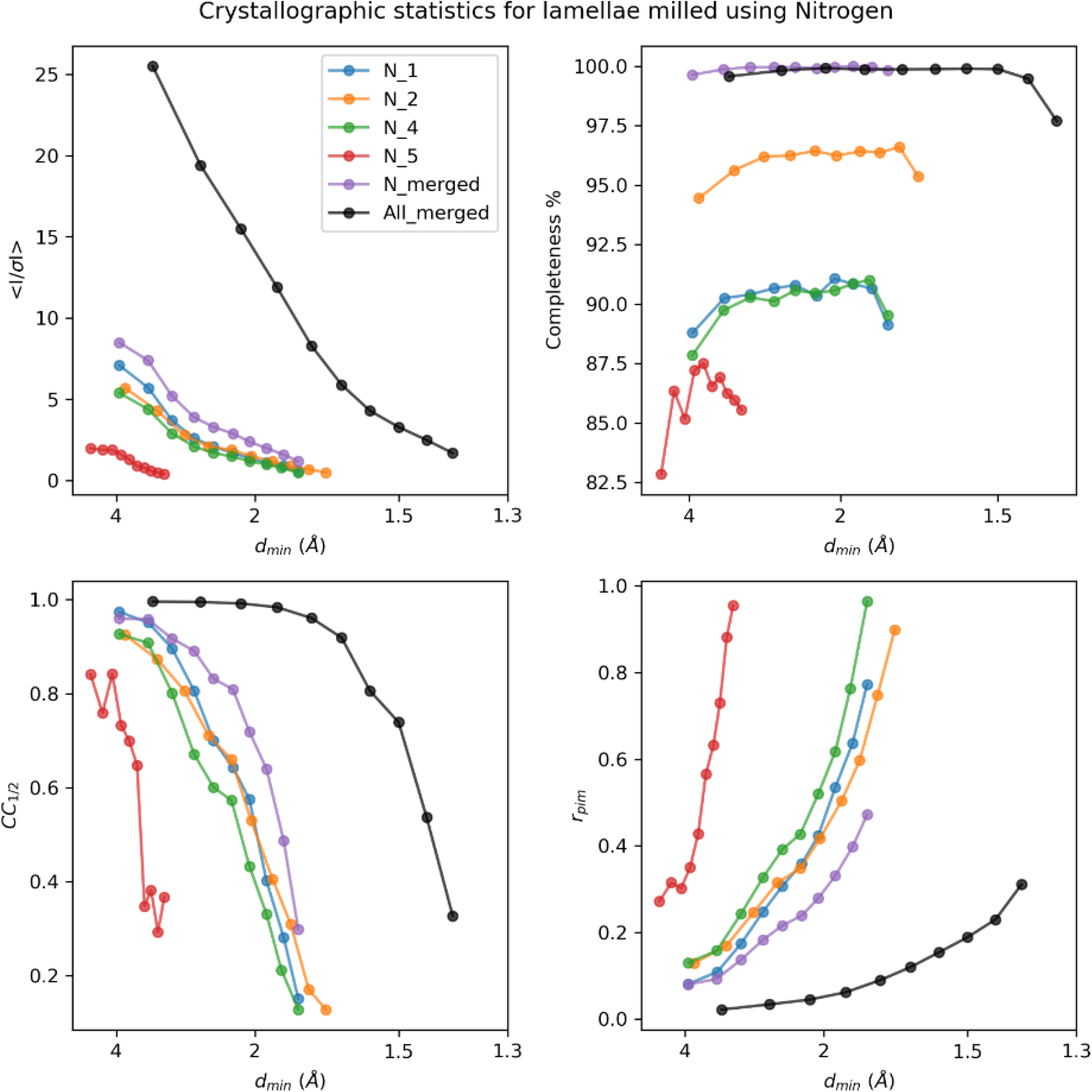
Crystallographic statistics for nitrogen ion-beam milled lamellae. Plots depict the mean signal to noise ratio (<I / σ (I)>) (top left), completeness (%) (top right), mean half-set correlation coefficient (CC_1/2_) (bottom left), and multiplicity corrected R factor (Rpim)(bottom right) as functions of the dmin resolution bins (Å). The “best merge” data set is included for comparison in each case.

**Supplemental Figure 14.**
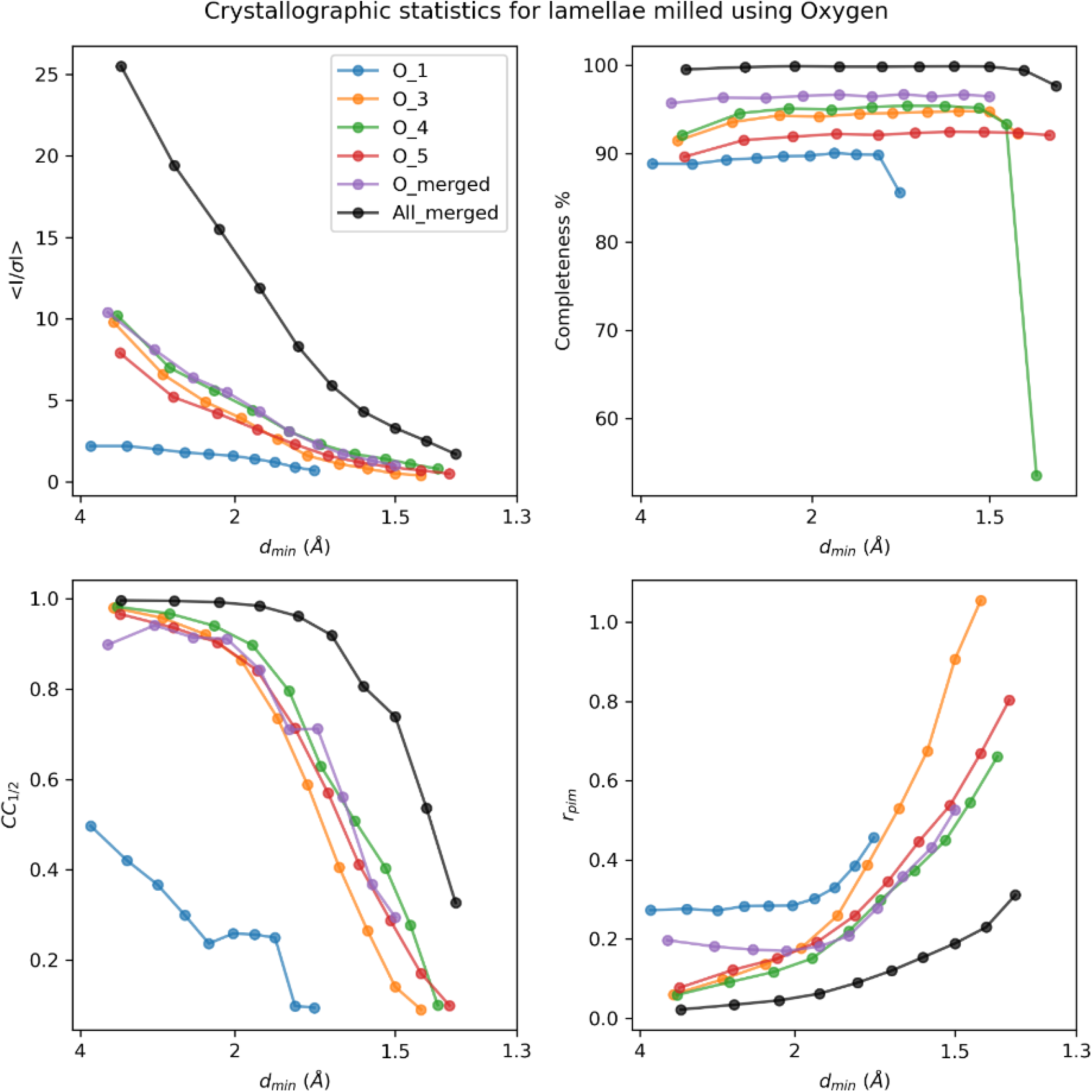
Crystallographic statistics for oxygen ion-beam milled lamellae. Plots depict the mean signal to noise ratio (<I / σ (I)>) (top left), completeness (%) (top right), mean half-set correlation coefficient (CC_1/2_) (bottom left), and multiplicity corrected R factor (Rpim)(bottom right) as functions of the dmin resolution bins (Å). The “best merge” data set is included for comparison in each case.

**Supplemental Figure 15.**
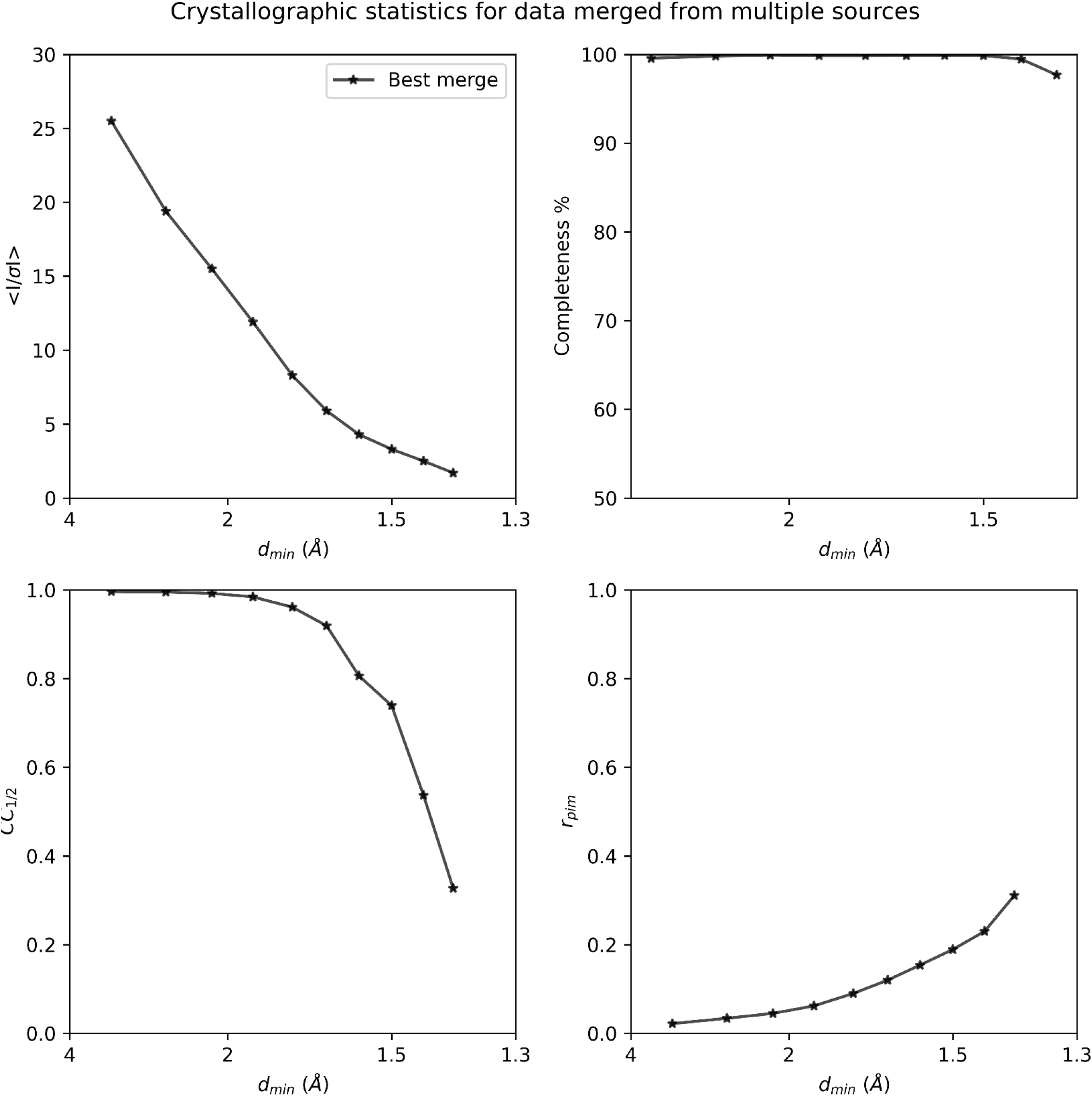
Crystallographic statistics for best merge from all ion-beam milled lamellae. Plots depict the mean signal to noise ratio (<I / σ (I)>) (top left), completeness (%) (top right), mean half-set correlation coefficient (CC_1/2_) (bottom left), and multiplicity corrected R factor (R_pim_)(bottom right) as functions of the dmin resolution bins (Å).

**Supplementary Table 1.** Milling steps for each plasma ion beam experiment on proteinase K.

| Xenon | Ion-beam current | Box size*<br>(x, y, z um) | Pattern type | Time for pattern x2 | Pattern separation |
| --- | --- | --- | --- | --- | --- |
| Step 1 | 1.0 nA | 6 x 6 x 5 | Cleaning cross section (CCS) | 5:28 | 2 um |
| Step 2 | 0.3 nA | 6 x 1 x 3 | CCS | 1:52 | 1 um |
| Step 3 | 0.1 nA | 5 x 0.5 x 2 | CCS | 1:34 | 500 nm |
| Step 4 | 30 pA | 5 x 0.5 x 1 | CCS | 2:36 | 300 nm |
| Total time milling patterns: |  |  |  | 11:30 |  |
| Argon | Ion-beam current | Box size*<br>(x, y, z um) | Pattern type | Time for pattern x2 | Pattern separation |
| Step 1 | 2.0 nA | 6 x 6 x 6 | Cleaning cross section (CCS) | 3:16 | 2 um |
| Step 2 | 0.74 nA | 6 x 1 x 3 | CCS | 0:46 | 1 um |
| Step 3 | 0.2 nA | 5 x 0.5 x 2 | CCS | 0:48 | 500 nm |
| Step 4 | 60 pA | 5 x 0.5 x 2 | CCS | 2:36 | 300 nm |
| Total time milling patterns: |  |  |  | 7:26 |  |
| Nitrogen | Ion-beam current | Box size*<br>(x, y, z um) | Pattern type | Time for pattern x2 | Pattern separation |
| Step 1 | 2.4 nA | 6 x 6 x 20 | Cleaning cross section (CCS) | 9:12 | 2 um |
| Step 2 | 0.78 nA | 5 x 1 x 10 | CCS | 2:04 | 1 um |
| Step 3 | 0.27 nA | 5 x 0.5 x 5 | CCS | 1:32 | 500 nm |
| Step 4 | 47 pA | 5 x 0.5 x 5 | CCS | 8:22 | 300 nm |
| Total time milling patterns: |  |  |  | 21:20 |  |
| Oxygen | Ion-beam current | Box size*<br>(x, y, z um) | Pattern type | Time for pattern x2 | Pattern separation |
| Step 1 | 1.7 nA | 6 x 6 x 15 | Cleaning cross section (CCS) | 8:14 | 2 um |
| Step 2 | 0.61 nA | 5 x 1 x 6 | CCS | 1:06 | 1 um |
| Step 3 | 0.23 nA | 5 x 0.5 x 4 | CCS | 1:06 | 500 nm |
| Step 4 | 90 pA | 5 x 0.5 x 4 | CCS | 3:40 | 300 nm |
| Total time milling patterns: |  |  |  | 14:06 |  |
| Gallium | Ion-beam current | Box size*<br>(x, y, z um) | Pattern type | Time for pattern x2 | Pattern separation |
| Step 1 | 0.5 nA | 6 x 6 x 10 | Cleaning cross section (CCS) | 7:04 | 2 um |
| Step 2 | 0.3 nA | 6 x 1 x 6 | CCS | 1:46 | 1 um |
| Step 3 | 0.1 nA | 5 x 0.5 x 3 | CCS | 1:06 | 500 nm |
| Step 4 | 30 pA | 5 x 0.5 x 3 | CCS | 3:30 | 300 nm |
| Total time milling patterns: |  |  |  | 13:26 |  |

**Supplementary Table 2.** Milling currents for each available ion source on the pFIB.

| Source | Xenon | Argon | Nitrogen | Oxygen |
| --- | --- | --- | --- | --- |
| Aperture #1 | 1pA | 3.9pA | 0.58pA | 1.3pA |
| #2 | 3.0pA | 6.0pA | 0.64pA | 1.4pA |
| #3 | 10pA | 20pA | 1.9pA | 4.2pA |
| #4 | 30pA | 60pA | 10pA | 20pA |
| #5 | 0.1nA | 0.2nA | 47pA | 90pA |
| #6 | 0.3nA | 0.74nA | 0.1nA | 0.23nA |
| #7 | 1.0nA | 2.0nA | 0.27nA | 0.61nA |
| #8 | 4.0nA | 7.6nA | 0.78nA | 1.7nA |
| #9 | 15nA | 28nA | 2.4nA | 5.6nA |
| #10 | 60nA | 0.12uA | 23nA | 45nA |
| #11 | 0.2uA | 0.40uA | 0.1uA | 0.19uA |
| #12 | 0.5uA | 0.93uA | 0.33uA | 0.57uA |
| #13 | 1.0uA | 2.0uA | 0.7uA | 1.0uA |
| #14 | 2.5uA | 4.0uA | 1.0uA | 2.0uA |

